# Glutamate dehydrogenase (Gdh2)-dependent alkalization is dispensable for escape from macrophages and virulence of *Candida albicans*

**DOI:** 10.1101/2020.01.20.912592

**Authors:** Fitz Gerald S. Silao, Kicki Ryman, Tong Jiang, Meliza Ward, Nicolas Hansmann, Ning-Ning Liu, Changbin Chen, Per O. Ljungdahl

## Abstract

*Candida albicans* cells depend on the energy derived from amino acid catabolism to induce and sustain hyphal growth inside phagosomes of engulfing macrophages. The concomitant deamination of amino acids is thought to neutralize the acidic microenvironment of phagosomes, a presumed requisite for survival and initiation of hyphal growth. Here, in contrast to an existing model, we show that mitochondrial-localized NAD^+^-dependent glutamate dehydrogenase (*GDH2*) catalyzing the deamination of glutamate to α-ketoglutarate, and not the cytosolic urea amidolyase (*DUR1,2*), accounts for the observed alkalization of media when amino acids are the sole sources of carbon and nitrogen. *C. albicans* strains lacking *GDH2* (*gdh2*-/-) are viable and do not extrude ammonia on amino acid-based media. Environmental alkalization does not occur under conditions of high glucose (2%), a finding attributable to glucose-repression of *GDH2* expression and mitochondrial function. Consistently, inhibition of oxidative phosphorylation or mitochondrial translation by antimycin A or chloramphenicol, respectively, prevents alkalization. *GDH2* expression and mitochondrial function are derepressed as glucose levels are lowered from 2% (∼110 mM) to 0.2% (∼11 mM), or when glycerol is used as carbon source. Using time-lapse microscopy, we document that *gdh2*-/- cells survive, filament and escape from primary murine macrophages at rates indistinguishable from wildtype. Consistently, *gdh2*-/- strains are as virulent as wildtype in fly and murine models of systemic candidiasis. Thus, although Gdh2 has a critical role in central nitrogen metabolism, Gdh2-catalyzed deamination of glutamate is surprisingly dispensable for escape from macrophages and virulence, demonstrating that amino acid-dependent alkalization is not essential for hyphal growth, survival in macrophages and hosts. An accurate description of the microenvironment within the phagosomal compartment of macrophages and the metabolic events underlying the survival of phagocytosed *C. albicans* cells and their escape are critical to understanding the host-pathogen interactions that ultimately determine the pathogenic outcome.

**Author Summary:** *Candida albicans* is a commensal component of the human microflora and the most common fungal pathogen. The incidence of candidiasis is low in healthy populations. Consequently, environmental factors, such as interactions with innate immune cells, play critical roles. Macrophages provide the first line of defense and rapidly internalize *C. albicans* cells within specialized intracellular compartments called phagosomes. The microenvironment within phagosomes is dynamic and ill defined, but has a low pH, and contains potent hydrolytic enzymes and oxidative stressors. Despite the inhospitable conditions, phagocytized *C. albicans* cells catabolize amino acids to obtain energy to survive and grow. Here, we have critically examined amino acid catabolism and ammonia extrusion in *C. albicans*, the latter is thought to neutralize the phagosomal pH and induce the switch of morphologies from round “yeast-like” to elongated hyphal cells that can pierce the phagosomal membrane leading to escape from macrophages. We report that Gdh2, which catalyzes the deamination of glutamate to α-ketoglutarate, is responsible for the observed environmental alkalization when *C. albicans* catabolize amino acids. Strikingly, Gdh2 is dispensable for escape from macrophages and virulent growth. These results provide new insights into host-pathogen interactions that determine the pathogenic outcome of *C. albicans* infections.

## INTRODUCTION

*Candida albicans* is a benign member of mucosal microbiota of most humans. However, in individuals with an impaired immune response, *C. albicans* can cause serious systemic infections associated with high rates of mortality (1, 2). In establishing virulent infections, *C. albicans* cells overcome potential obstacles inherent to the microenvironments in the host. Consistently, the capacity of *C. albicans* to establish a wide spectrum of pathologies is attributed to multiple virulence factors, one of which involves morphological switching from the yeast to filamentous forms (i.e., hyphae and pseudohyphae), reviewed in (3–5). The ability to switch from yeast to filamentous growth is required for tissue invasion and escape from innate immune cells, such as macrophages, whereas, the yeast form facilitates dissemination via the bloodstream. In addition to escaping from innate immune cells, fungal cells must successfully compete with host cells and even other constituents of the microbiome to take up necessary nutrients for growth (6).

Phagocytes, such as macrophages, are among the first line of host defenses encountered by *C. albicans* (reviewed in (7)). These innate immune cells recognize specific fungal surface antigens via specific plasma membrane-bound receptors (8). Once recognized, fungal cells are enveloped by membrane protrusions that form the phagosomal compartment. The phagosome matures by fusing with discrete intracellular organelles, resulting in a compartment with potent hydrolytic enzymes, oxidative stressors and low pH (8–10). Acidification is important to optimize the activity of the hydrolytic enzymes that target to the phagosome.

*C. albicans* can survive and even escape macrophage engulfment. This is thought to depend on the ability of fungal cells to raise the phagosomal pH via ammonia extrusion. It has been suggested that urea amidolyase (Dur1,2), localized to the cytoplasm, catalyzes the reactions generating the ammonia extruded from cells by the plasma membrane-localized Ato proteins (11, 12). In addition to impairing the activity of pH-sensitive proteolytic enzymes, phagosomal alkalization is thought to initiate and promote hyphal growth (11, 13). Consistent with this notion, *C. albicans* lacking *STP2*, encoding one of the SPS (Ssy1-Ptr3-Ssy5) sensor controlled effector factors governing amino acid permease gene transcription (14), fail to form hyphae and escape macrophages (13). These observations led to a model that the reduced capacity of *stp2*Δ strains to take up amino acids limits the supply of substrates of Dur1,2 catalyzed deamination reactions, which would result in the reduced capacity to alkalinize the phagosome (12, 13).

We have recently shown that the mitochondrial proline catabolism is required for hyphal growth and macrophage evasion. The proline catabolic pathway is the primary route of arginine utilization (15) and operates independently of the cytosolic Dur1,2-catalyzed urea-CO_2_ pathway (15, 16). In contrast to the proposed model (12), we observed that *dur1,2-/-* cells retain the capacity to alkalinize a basal medium containing arginine as sole nitrogen and carbon source (15). Furthermore, strains carrying *put1*-/- or *put2*-/- mutations exhibit strong growth defects and consequently, are incapable of alkalinizing the same medium, suggesting that alkalization is linked to proline catabolism.

A potential source of ammonia responsible for alkalization is the deamination of glutamate, a metabolic step downstream of Put2. In *Saccharomyces cerevisiae*, the primary source of free ammonia is generated by the mitochondrial-localized NAD^+^- dependent glutamate dehydrogenase (Gdh2) catalyzed deamination of glutamate to α-ketoglutarate, a reaction that generates NADH and NH3 (17). Importantly, the reaction is anaplerotic and replenishes the tricarboxylic acid (TCA) cycle with α-ketoglutarate, a key TCA cycle intermediate between isocitrate and succinyl CoA, and an important precursor for amino acid biosynthesis.

Here, we have examined the role of Gdh2 in morphological switching under *in vitro* conditions in filament-inducing media, *in situ* in the phagosome of primary murine macrophages, and in virulence in two model host systems. We show that when *C. albicans* utilize amino acids as sole nitrogen- and carbon-sources they extrude ammonia, which originates from Gdh2-catalyzed deamination of glutamate. In contrast to current understanding regarding the importance of phagosomal alkalization, we report that *C. albicans* strains lacking *GDH2* filament and escape the phagosome of engulfing macrophages at rates indistinguishable to wildtype. Furthermore, we also report that the Gdh2-catalyzed reaction is dispensable for virulence in both fly and murine models of systemic candidiasis.

## RESULTS

### *C. albicans GDH2* is responsible for amino acid-dependent alkalization *in vitro*

Arginine is rapidly converted to proline and then catabolized to glutamate in the mitochondria through the concerted action of two enzymes, proline oxidase (Put1; proline to Δ^1^-pyrroline-5-carboxylate or P5C) and P5C dehydrogenase (Put2; P5C to glutamate) (**Fig. 1A**). *C. albicans* strains lacking *PUT1* (*put1*-/-) and/or *PUT2* (*put2*-/-) are unable to grow efficiently in minimal medium containing 10 mM of arginine as sole nitrogen and carbon source (YNB+Arg, pH = 4.0), and fail to alkalinize the medium (15). In contrast, cells carrying null alleles of *DUR1,2* (*dur1,2*-/-) grow robustly and alkalinize the media (15). To test if the catabolism of amino acids other than arginine and proline can be used as sole carbon source we examined the growth characteristics of the strains in YNB containing 1% casamino acids, a medium containing high levels of all proteinogenic amino acids (**Fig. 1B**). In this media, *dur1,2*-/- cells grew as wildtype and readily alkalinized the media (compare tube 5 with 6). In contrast, *put1-/-* cells exhibited poor growth and weakly alkalinized the medium (tube 3). Cells lacking Put2 activity (*put2-/-*) did not grow and the culture media remained acidic (tube 2). Interestingly, a *put1-/- put2-/-* double mutant strain grew better than the single *put2-/-* mutant (compare tube 4 with 2). The severe growth impairment associated with the loss of Put2 is likely due to the accumulation of P5C, which is known to cause mitochondrial disfunction (18). These results indicate that the amino acids metabolized via the proline catabolic pathway are preferentially used as carbon sources when mixtures of amino acids are present, e.g., in casamino acid preparations. The catabolism of these non-prefered amino acids contribute only modestly to alkalization, consistent with reports that not all amino acids can serve as carbon sources and contribute to environmental alkalization (12).

**Figure 1.**
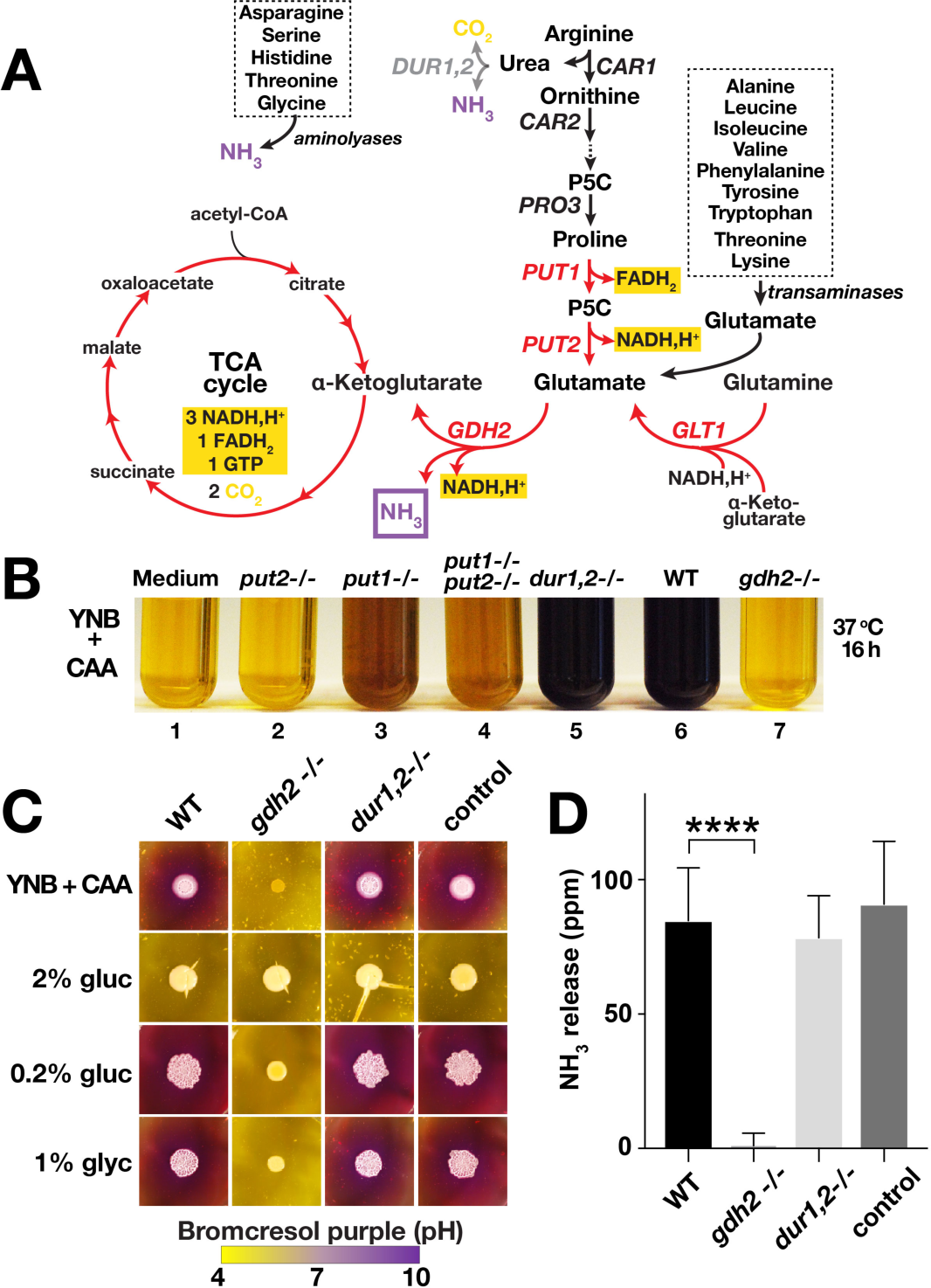
*C. albicans GDH2* is required for growth using amino acids as sole carbon and nitrogen sources. (A) Schematic diagram of arginine/proline catabolism. Arginine is catabolized to proline in the cytoplasm, proline is transported into mitochondria, proline is catabolized to glutamate in two enzymatic reactions, catalyzed by FAD-dependent proline oxidase (*PUT1*) and NAD^+^-linked Δ^1^-pyrroline-5-carboxylate (P5C) dehydrogenase (*PUT2*), respectively. The two central reactions for nitrogen source utilization are catalyzed by NADH-dependent glutamate synthase (*GLT1*) and NAD^+^-linked glutamate dehydrogenase (*GDH2*). The gene products and metabolic steps marked in red are localized to the mitochondria. (B) YPD grown *put2-/-* (CFG318), *put1*-/- (CFG154), *put1-/- put2-/-* (CFG159), *dur1,2-/-* (CFG246), wildtype (WT, SC5314) and *gdh2-/-* (CFG279) cells were washed, resuspended at an OD_600_ of 0.05 in YNB+CAA containing the pH indicator bromocresol purple, and the cultures were incubated shaking at 37 °C for 16 h. (C) Wildtype (WT, SC5314), *gdh2-/-* (CFG279), *dur1,2-/-* (CFG246) and CRISPR control (CFG182) cells were pre-grown in YPD, washed, resuspended at an OD_600_ of 1, and 5 μl were spotted onto the surface of solid YNB + CAA bromocresol purple without and with the indicated carbon source. The plates were incubated at 37 °C for 72 h. The images are representative of at least 3 independent experiments. (D) Volatile ammonia released from strains as in (C); the results are the average of at least 3 independent experiments (Ave. ± CI; **** p ≤ 0.0001).

The requirement of proline catabolism for growth suggested that the downstream deamination of glutamate to α-ketoglutarate, catalyzed by glutamate dehydrogenase, provided the metabolite responsible for alkalinizing the media. In *S. cerevisiae*, mitochondrial glutamate dehydrogenase (*GDH2*) is the primary source of free ammonia (17). The *C. albicans* genome has one gene predicted to encode glutamate dehydrogenase (*GDH2*, C5_02600W), and using CRISPR/Cas9 we inactivated both alleles of this gene in SC5314. Two independent clones were analyzed (**Fig. S1A and S1B)**. The *gdh2-/-* strains were viable on YPD or YPG (**Fig. S1C**), however, they showed strong growth and alkalization defects when amino acids were used as sole nitrogen and carbon sources, such as in YNB+Arg (**Fig. S1A**) and YNB+CAA media (**Fig.1B, 1C, and S2A**). Consistent with what is known for *S. cerevisiae*, the *gdh2*-/- mutant showed a modest growth defect in media containing glutamate or proline as sole nitrogen source (**Fig. S1C**).

To further test whether Gdh2 is responsible for environmental alkalization we assessed the capacity of the *gdh2*-/-mutant to grow and alkalinize the external growth milieu on solid YNB+CAA (**Fig. 1C**). In the absence of an additional carbon source, cells lacking *GDH2* did not grow appreciably, and failed to form colonies. By contrast, on YNB+CAA supplemented with 2% glucose the *gdh2-/-* strain formed colonies of similar size as WT, indicating that Gdh2 is dispensable for growth in media with high levels of glucose. Consistent with a requirement of a fermentable carbon source, the *gdh2-/-* strain exhibited reduced growth on media with 0.2% glucose, or the non-fermentable carbon source glycerol (1%). Although able to grow in the presence of an added carbon source, the *gdh2-/-* stain failed to alkalinize the media. In contrast, wildtype (WT), *dur1,2-/-* and CRISPR control cells formed colonies of equal size on all media and exhibited identical capacities to alkalinize the media (**Fig. 1C**, compare columns 3 and 4 with 1). These observations confirm that Gdh2 is responsible for alkalization of the external growth environment.

### *GDH2* is required for ammonia extrusion

Next, we analyzed whether the alkalization defect of the *gdh2*-/- mutant was due to the lack of ammonia extrusion. The levels of volatile ammonia produced was measured by colonies growing on solid YNB+CAA with 0.2% glucose medium buffered with MOPS (pH = 7.4); the standard acidic growth medium (pH = 4.0) traps ammonia (NH^3^) as ammonium (NH_4_ ^+^), decreasing the level of volatile ammonia and thereby interfering with the assay. As shown in **Fig. 1D**, the *gdh2*-/- strain did not release measurable ammonia. Consistent with their ability to alkalinize the growth media (**Fig. 1C**), wildtype, *dur1,2*-/- and CRISPR control strains released substantial and indistinguishable levels of ammonia. Together, these results indicate that the reaction catalyzed by Gdh2 generates the ammonia that alkalinizes the growth environment when *C. albicans* uses amino acids as the primary energy source.

### Environmental alkalization originates in the mitochondria

We recently confirmed that mitochondrial activity in *C. albicans* can be repressed by glucose (15), a finding that is consistent with existing transcriptional profiling data (19). Consequently, the glucose repressible nature of extracellular alkalization in the presence of amino acids could be linked to glucose repressed mitochondrial function. To examine this notion, we first sought to confirm that Gdh2 localizes to mitochondria. Cells (CFG273) expressing the functional *GDH2-GFP* reporter were grown in synthetic glutamate media with 0.2% glucose (SED_0.2%_) and YNB+CAA. The GFP fluorescence in cells grown under both conditions clearly localized to the mitochondria as determined by the precise overlapping pattern of fluorescence with the mitochondrial marker MitoTracker Deep Red (MTR) (**Fig. 2A**).

**Figure 2.**
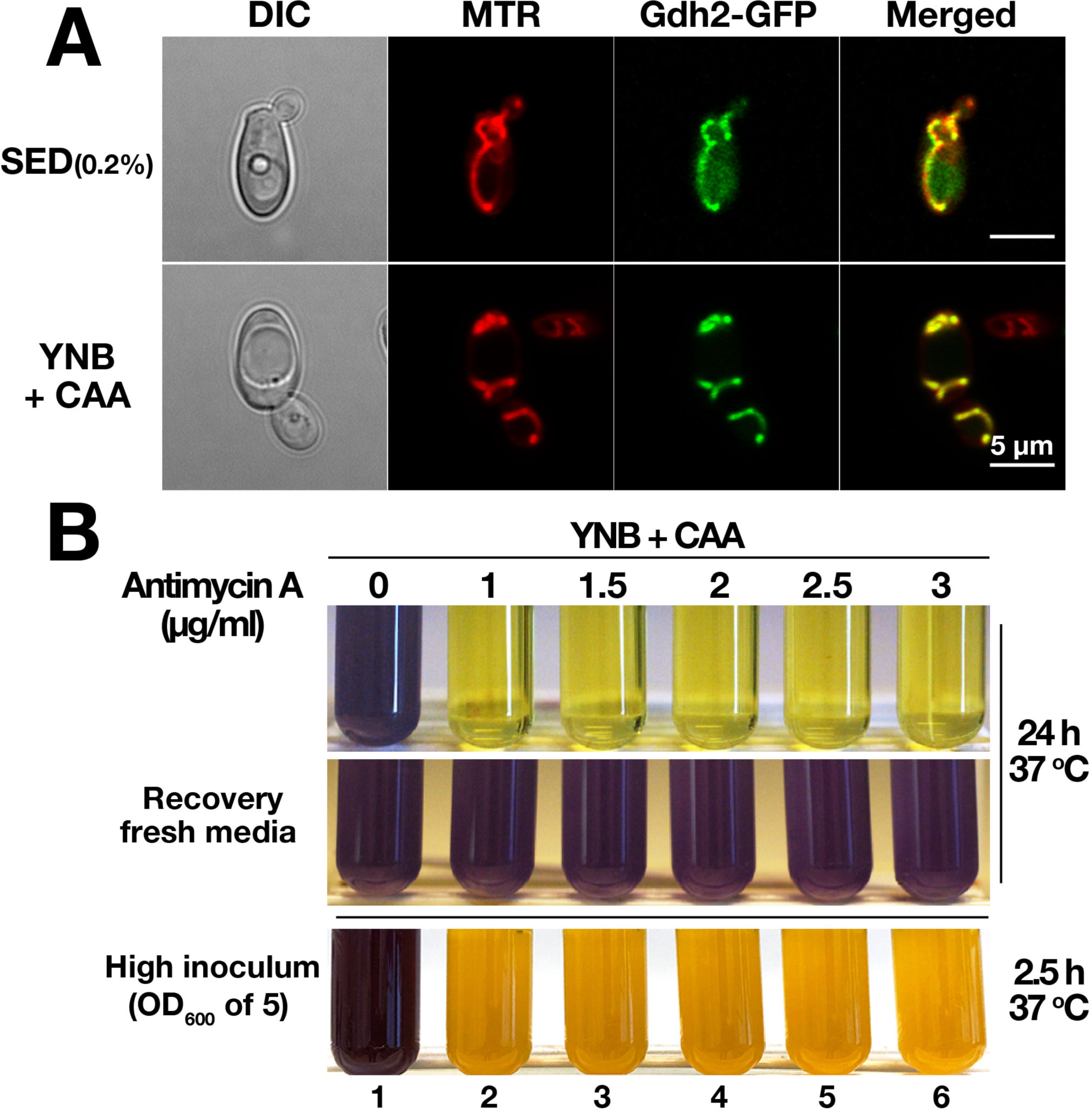
*C. albicans* Gdh2 localizes to the mitochondria and environmental alkalization requires mitochondrial function. (A) Gdh2-GFP co-localizes with the mitochondrial marker MitoTracker Red (MTR). YPD grown cells expressing *GDH2-GFP* (CFG273) were harvested, washed, grown in SED (0.2% glucose) or YNB + CAA at 37 °C for 24 h, and stained with 200 nM MTR prior to imaging by differential interference contrast (DIC) and confocal fluorescence microscopy; the scale bar = 5 μm. (B) Wildtype cells (SC5314) from overnight YPD cultures were washed and then diluted to either OD_600_ ≈ 0.1 (top panel) or ≈ 5 (bottom panel) in liquid YNB+CAA with the indicated concentrations of mitochondrial complex III inhibitor antimycin A. Cultures were grown at 37 °C under constant aeration for 24 h and 2.5 h, respectively, and photographed. To assess viability after Antimycin A treatment, inhibited cells from 24 h old culture (top panel) were harvested, washed, and then resuspended in fresh YNB+CAA media and incubated for 24 h (up to 48 h) at 37 °C (middle panel). Images are representative of at least 3 independent experiments.

To independently assess the role of mitochondrial activity in the alkalization process, we grew the wildtype cells in standard YNB+CAA medium (without glucose), in the presence of Antimycin A, a potent inhibitor of respiratory complex III. No alkalization was observed in the medium even after 24 h of growth (**Fig. 2B**, upper left panel). Antimycin A clearly impeded the growth of wildtype cells, which phenocopies the *gdh2*-/- growth in YNB+CAA. To ascertain whether the failure to alkalinize the medium was due to inhibiting mitochondrial respiration and not due to cell death, we harvested the cells from antimycin-treated cultures and suspended them in fresh medium; the cells regained their capacity to alkalinize the medium (**Fig. 2B**, lower left panel). To further test that the inhibitory effect of antimycin A on alkalization is specific to mitochondrial function and not an indirect effect of growth inhibition, we performed the same experiment but starting with a high cell density (OD_600_ of 5). As shown in **Fig. 2B** (right panel), the color endpoint indicating alkalization in wildtype control occurred in a span of 2.5 h following inoculation whereas all antimycin A-treated cells failed to neutralize the pH of the medium. This clearly demonstrates that mitochondrial function is essential for environmental alkalization. We also grew the cells in the presence of chloramphenicol, a potent mitochondrial inhibitor that targets mitochondrial translation by reversibly binding to the 50S subunit of the 70S ribosome in yeast (20). In the presence of this inhibitor and a low starting cell density, a delay in alkalization was observed initially (**Fig. S2A**), but then alkalization was virtually indistinguishable after 48 h, or when a high starting cell density was used (data not shown). These results, together with our observation that glucose availability influences Gdh2-dependent growth and alkalization (**Fig. 1C**), support our conclusion that alkalization originates from metabolism localized to mitochondria.

### Gdh2 expression is repressed by glucose

To follow up on the observations that glucose negatively affects Gdh2 activity and Gdh2 is a component of mitochondria (**Fig. 2A**), we sought to visualize Gdh2 expression in living cells when shifted from repressing YPD (2% glucose) to non-repressing YNB+CAA. To do this, we used the same Gdh2-GFP reporter strain described earlier (**Fig. 2A**). This enabled us to observe Gdh2 expression in single cells growing on a thin YNB+CAA agar slab over a period of 6 h. The Gdh2-GFP signal was initially weak (t = 0 h), becoming more intense as time progressed and as cells underwent several rounds of cell division, some leading to filamentous pseudohyphal growth (**Fig. 3A**).

**Figure 3.**
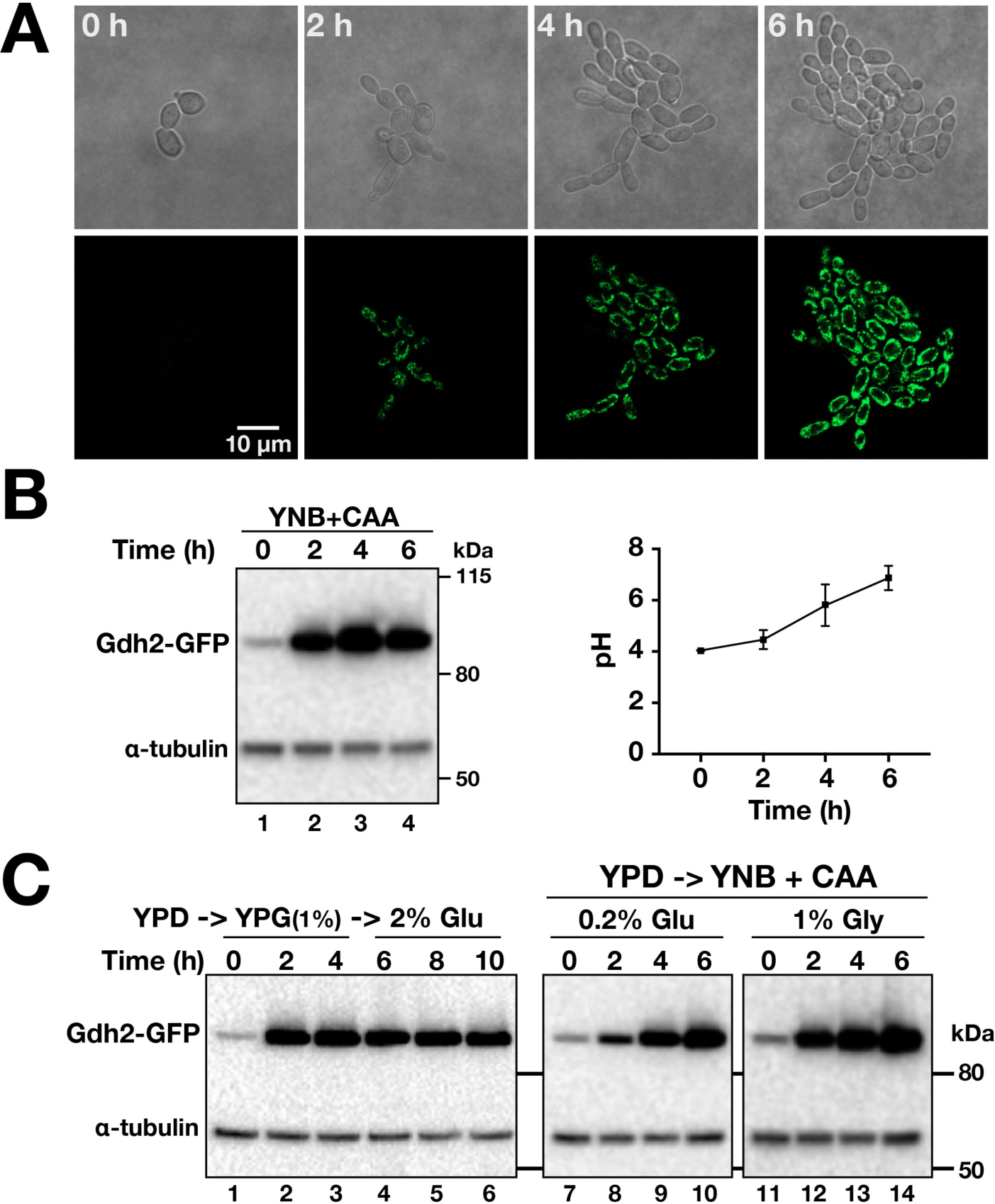
*GDH2* expression is repressed by glucose. (A) Live cell imaging of Gdh2-GFP expression in cells shifted from YPD to YNB+CAA. CFG273 (Gdh2-GFP) cells were pre-grown in YPD and transferred to a thin agar slab of YNB+CAA medium. Growth at 37 °C was monitored every hour for 6 h. (B) Gdh2-GFP expression is rapidly induced in cells shifted from YPD to YNB+CAA. Cells (same strain as in A) were pre-grown in YPD and used to inoculate liquid YNB+CAA (OD_600_ of 2.0); at the times indicated, the pH was measured (right panel; average of 3 independent experiments) and the levels of Gdh2-GFP expression (left panel) were monitored by immunoblot analysis. (C) Gdh2 expression is carbon source dependent. Cells (same strain as in A) grown in YPD were harvested, transferred to YPG (YP + 1% glycerol; lanes 1-3) and after subsampling at 6 hr, 2% glucose was added to cultures (lanes 5-6) (left panel); YPD grown cells were shifted to YNB + CAA with 0.2% glucose (lanes 7-10) or 1% glycerol (lanes 11-14) and grown at 37 °C (right panels). Extracts were prepared at the times indicated and the levels of Gdh2-GFP and tubulin (loading control) were assessed by immunoblotting using primary α-GFP and α-tubulin antibodies.

To relate this observation with the actual alkalization process, we analyzed the levels of Gdh2-GFP in cells grown in liquid culture taken at similar time points. Cells, pre-grown in YPD (2% glucose), were shifted to YNB+CAA and the levels of Gdh2-GFP were assessed by immunoblot analysis. To enable the recovery of adequate amounts of cells for subsequent extract preparation, we increased the starting cell density of the culture (i.e., OD_600_ ≈ 2.0). As shown in **Fig. 3B** (left panel) the Gdh2- GFP level in YPD-grown cells was initially low (t = 0 h) but within 2 h the level was greatly enhanced and remained so during the entire 6 hr incubation. During the course of growth, the media became successively alkaline, increasing from the starting pH of 4 to 7 (**Fig. 3B**, right panel). The finding that Gdh2 expression is induced in cells growing in media rich in amino acids (i.e., YNB+CAA or YPG) indicates that *GDH2* expression in *C. albicans*, in contrast to *S. cerevisiae* (21), is not subject to NCR.

Next, we examined the expression and stability of Gdh2-GFP in cells shifted from YPD to YPG (**Fig. 3C**). Again, the level of Gdh2-GFP rapidly increased (lanes 1-3) and remained high following the addition of glucose (2% final concentration) (**Fig. 3C**, lanes 5-6), an observation reminiscent of isocitrate lyase (Icl1), a glyoxylate cycle enzyme that is not subject to catabolite inactivation in *C. albicans* (22, 23). To further illustrate the effect of glucose on Gdh2-GFP expression, we shifted YPD grown cells to YNB+CAA in the presence of 0.2% glucose or 1% glycerol, conditions that are not repressive to mitochondrial function (15). As shown in **Fig. 3C**, the level of Gdh2-GFP substantially increased, clearly demonstrating that Gdh2 expression is sensitive to repression by glucose.

### Inactivation of Gdh2 does not impair morphogenesis

Based on current understanding that the ability of *C. albicans* to alkalinize their growth environments contributes to the induction of hyphal growth, and the observation that *gdh2*-/- cells formed smooth macrocolonies on YNB+CAA (pH = 4.0) with non-repressing 0.2% glucose and 1% glycerol (**Fig. 1C, column 2**), we examined whether the inactivation of Gdh2 would negatively affect morphogenesis. To test this notion, we examined growth on Spider and Lee’s media, two standard media used to assess filamentation. These media contain amino acids, have a neutral pH, and are known to promote filamentous growth of wildtype cells. Similar to wildtype and CRISPR control strains, the macrocolonies formed by the *gdh2-/*-strain were wrinkled and surrounded by an extensive outgrowth of hyphal cells (**Fig. 4A**). This indicates that Gdh2 function is dispensable for filamentation. This is supported by the recent paper showing the capacity of *gdh2*Δ/Δ to switch in amino acid-based medium (24).

**Figure 4.**
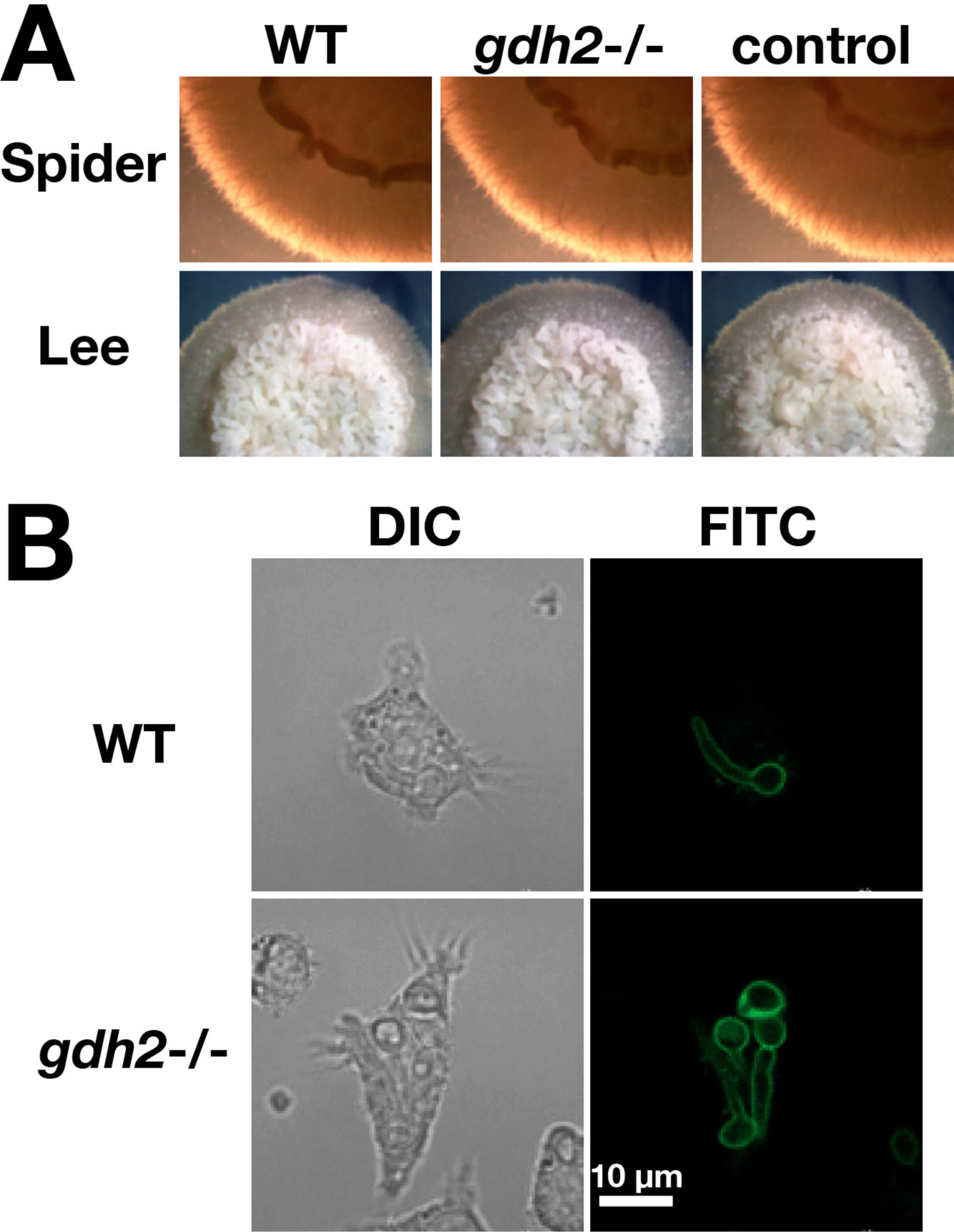
Gdh2 is dispensable for filamentous growth on solid media and inside phagosomes of engulfing macrophages. (A) Wildtype (WT, SC5314), *gdh2-/-* (CFG279), and CRISPR control (CFG182) strains, pre-grown in YPD, were washed, resuspended at an OD_600_ of 1 in water, and 5 μl aliquots were spotted on solid Spider and Lee’s media. Representative colonies were photographed 5 days after incubation at 37 °C. (B) FITC-stained WT (SC5314) and *gdh2*-/- (CFG279) cells were individually co-cultured with RAW264.7 macrophages. Non-phagocytosed fungal cells were removed by washing and the co-cultures were monitored by live cell imaging for 1.5 h; scale bar = 10 μm.

This unexpected result led us to evaluate the capacity of *gdh2*-/- cells to filament within phagosomes of engulfing macrophages. FITC-stained WT (SC5314) and *gdh2*-/- (CFG279) cells were individually co-cultured with RAW264.7 macrophages. Non-phagocytosed fungal cells were removed and then co-cultures were imaged after 1.5 h of incubation. We readily observed macrophages containing *gdh2-/-* cells that had formed hyphal extensions (**Fig. 4B**), clearly suggesting that amino acid-dependent alkalization of the phagosome is not a requisite for the induction of filamentous growth.

### Gdh2-GFP expression is rapidly induced upon phagocytosis by macrophages

To assess the time-course of Gdh2-GFP expression in phagocytized *C. albicans* cells, we co-cultured the Gdh2-GFP reporter strain CFG273 with RAW264.7 (RAW) macrophages and followed the interaction by time-lapse microscopy. The Gdh2- GFP signal significantly increased after phagocytosis (**Fig. 5A**, see Video V1, Supporting Information). We repeated the experiment using primary murine bone marrow-derived macrophage (BMDM) and obtained a similar result. However, due to the inherent green autofluorescence of BMDM, the GFP fluorescence appeared less pronounced (**Fig. 5B**, see Video V2, Supporting Information). These results demonstrate that although Gdh2 expression is induced rapidly following phagocytosis, presumably the reflection of limiting glucose availability and the subsequent release from glucose repression.

**Figure 5.**
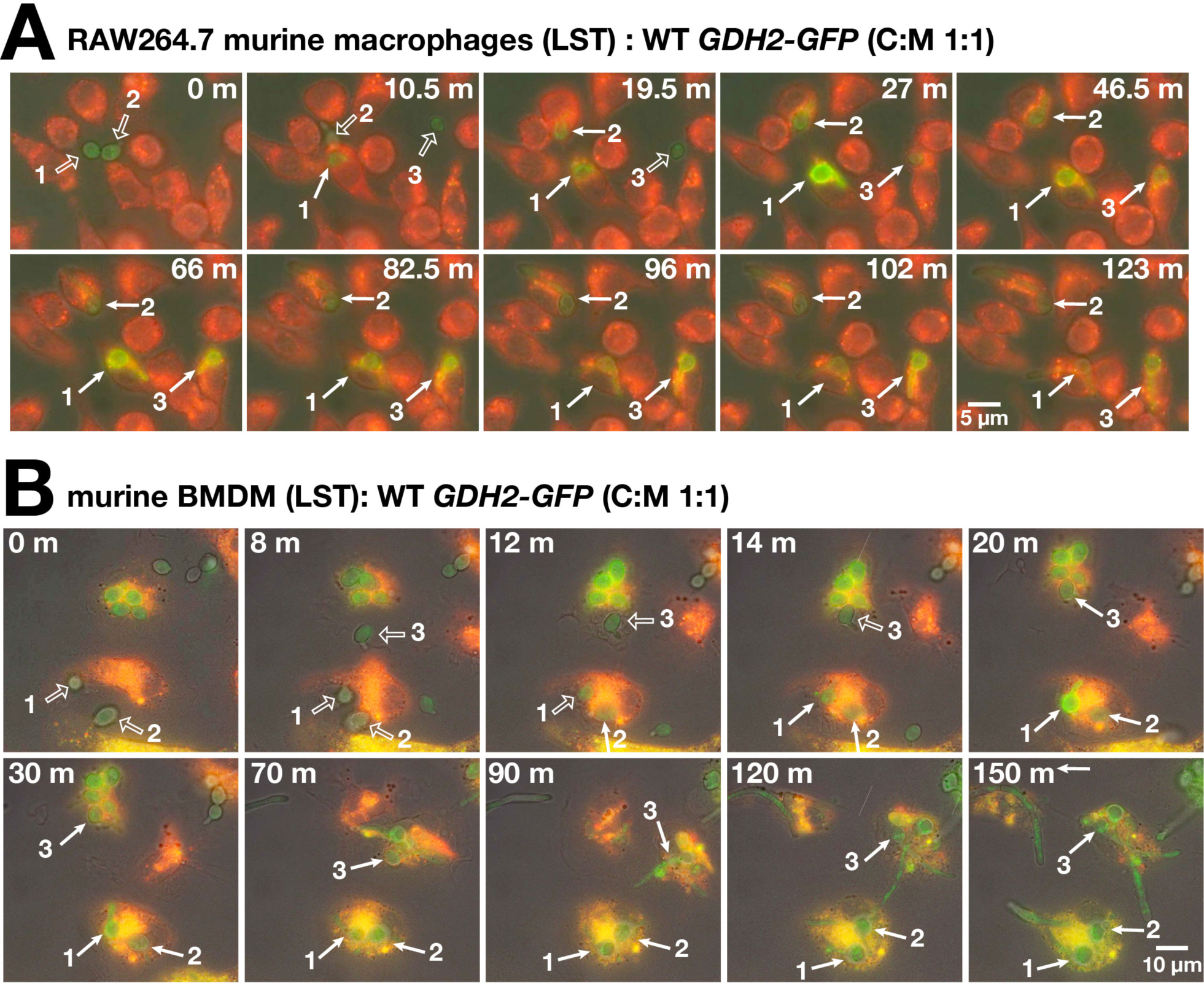
Gdh2-GFP expression is induced in *C. albicans* cells phagocytized by murine macrophages. CFG273 cells were co-cultured in CO_2_-independent medium with (A) RAW264.7 macrophages, or (B) primary murine bone marrow-derived macrophages (BMDM) pre-stained with Lysotracker Red (LST) at MOI of 1:1 (C:M). The co-cultures were followed by live cell imaging. Micrographs were taken at the times indicated (see Videos V1 and V2, Supporting Information). In each series, three CFG273 cells are marked prior to (open arrows) and after (closed arrows) being phagocytized.

### Gdh2 activity is not required to escape macrophages

We directly compared the ability of wildtype and the *gdh2*-/- mutant cells to survive and escape after being phagocytized by primary BMDM using a competition assay (**Fig. 6**). To carry out the experiment we created a wildtype strain constitutively expressing GFP (*ADH1*/*P_ADH1_*-GFP) and a *gdh2*-/- mutant strain constitutively expressing RFP (*gdh2*-/- *ADH1*/*P_ADH1_*-RFP). Both strains exhibited unaltered growth characteristics (see **Fig. S2B**). Equal numbers of WT and *gdh2-/-* cells were mixed (green:red; 1:1), and the fungal cell suspension was incubated with BMDM at a MOI of 3:1 in HBSS for 30 min before washing non-phagocytosed fungal cells away. The co-cultures were monitored by time-lapse microscopy.

**Figure 6.**
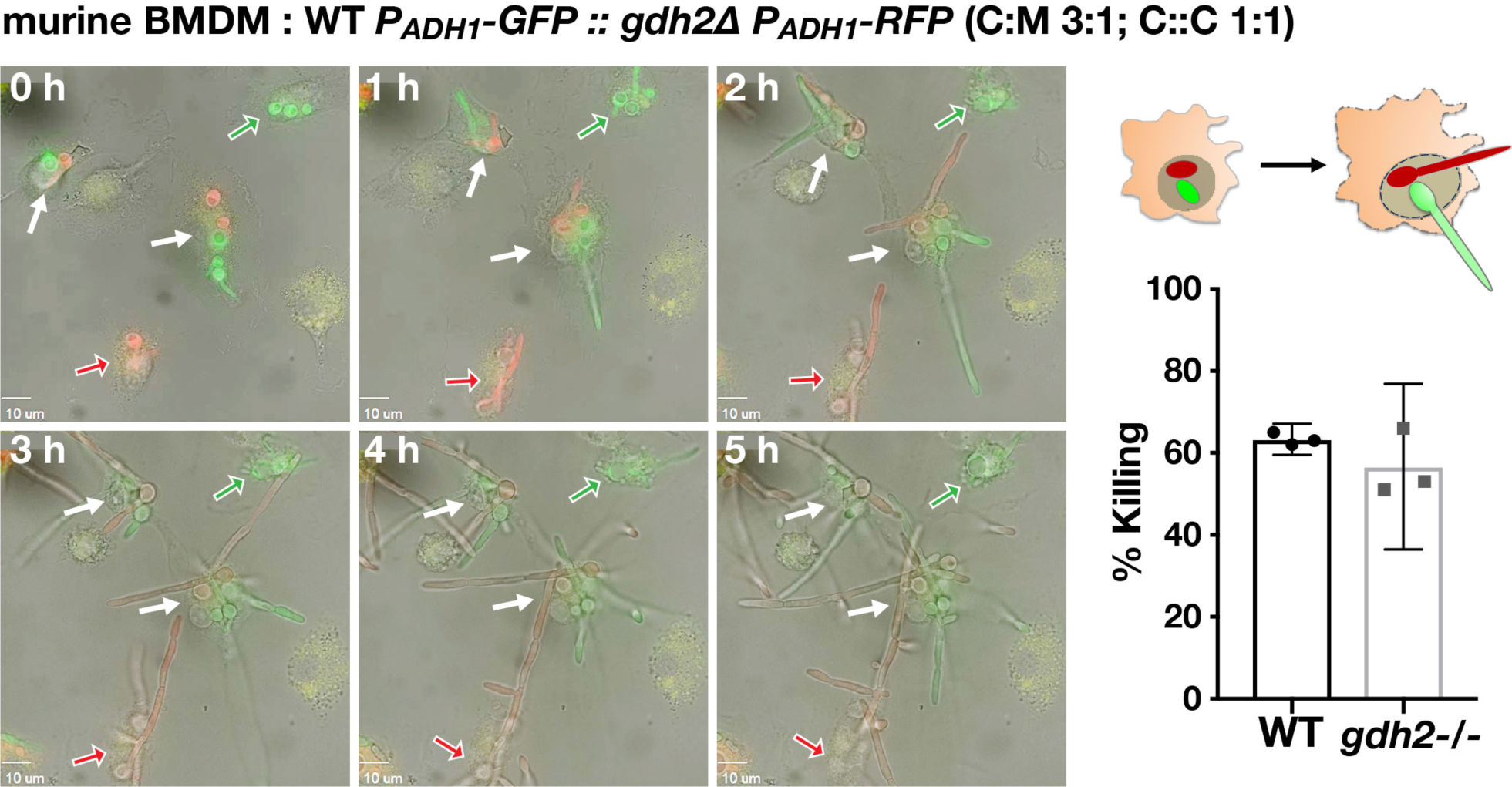
Competition assay to compare wildtype and *gdh2-/-* filamentation and survival upon phagocytosis by primary BMDM. Wildtype (WT; *P_ADH1_-GFP*; SCADH1G4A) and *gdh2*/- (*P_ADH1_-RFP*, CFG275) cells (1:1) were co-cultured with primary BMDM (MOI of 3:1; C:M) for 30 min in HBSS. Non-phagocytosed fungal cells were removed by washing and the co-culture was monitored by live cell imaging for 5 h (see video V3, Supporting Information). Solid arrows indicate macrophages with phagosomes containing both WT and *gdh2*-/- cells; open arrows indicate macrophages with phagosomes containing either WT (green) or *gdh2*-/- (red) cells. The observed growth of WT and *gdh2*-/- cells within a single macrophage is schematically illustrated (right upper panel). Candidacidal activity of BMDM (right lower panel); CFUs recovered at 2 h were compared to the CFUs in the starting inoculum.

Again, contrary to what we expected, the *gdh2*-/- mutant remained fully competent to initiate hyphal growth in the phagosome of BMDM (**Fig. 6**). Furthermore, given the perceived importance of environmental alkalization in the onset of phagosomal escape by *C. albicans*, we anticipated that the *gdh2*-/- mutant would be killed more efficiently than the wildtype. To test this notion, we performed a colony forming unit (CFU) assay to quantify the survival of phagocytized cells. The results show that Gdh2-mediated alkalization is not essential for survival in BMDM (**Fig. 6**, lower right panel), a result that is consistent to the recent report by Westman et al. (25). Together, our data indicate that ammonia extrusion and environmental alkalization are not requisites for the initiation of hyphal formation, growth and survival in the phagosome of primary BMDM.

### Gdh2 activity is dispensable for virulence in intact host

Next, we examined the role of Gdh2-dependent alkalization in the capacity of *C. albicans* to successfully infect an intact living host. We used an improved fruit fly (*Drosophila melanogaster*) infection model with the *Bom*^Δ*55C*^ flies that lack 10 Bomanin genes on chromosome 2 that encode for secreted peptides with antimicrobial property (26). As shown in the survival curve, the *gdh2*-/- mutant remained competent to infect *Bom*^Δ*55C*^ flies similar to wildtype control (**Fig. 7A**). The data indicate that Gdh2-dependent alkalization is not required for virulence in a *Drosophila* infection model.

**Figure 7.**
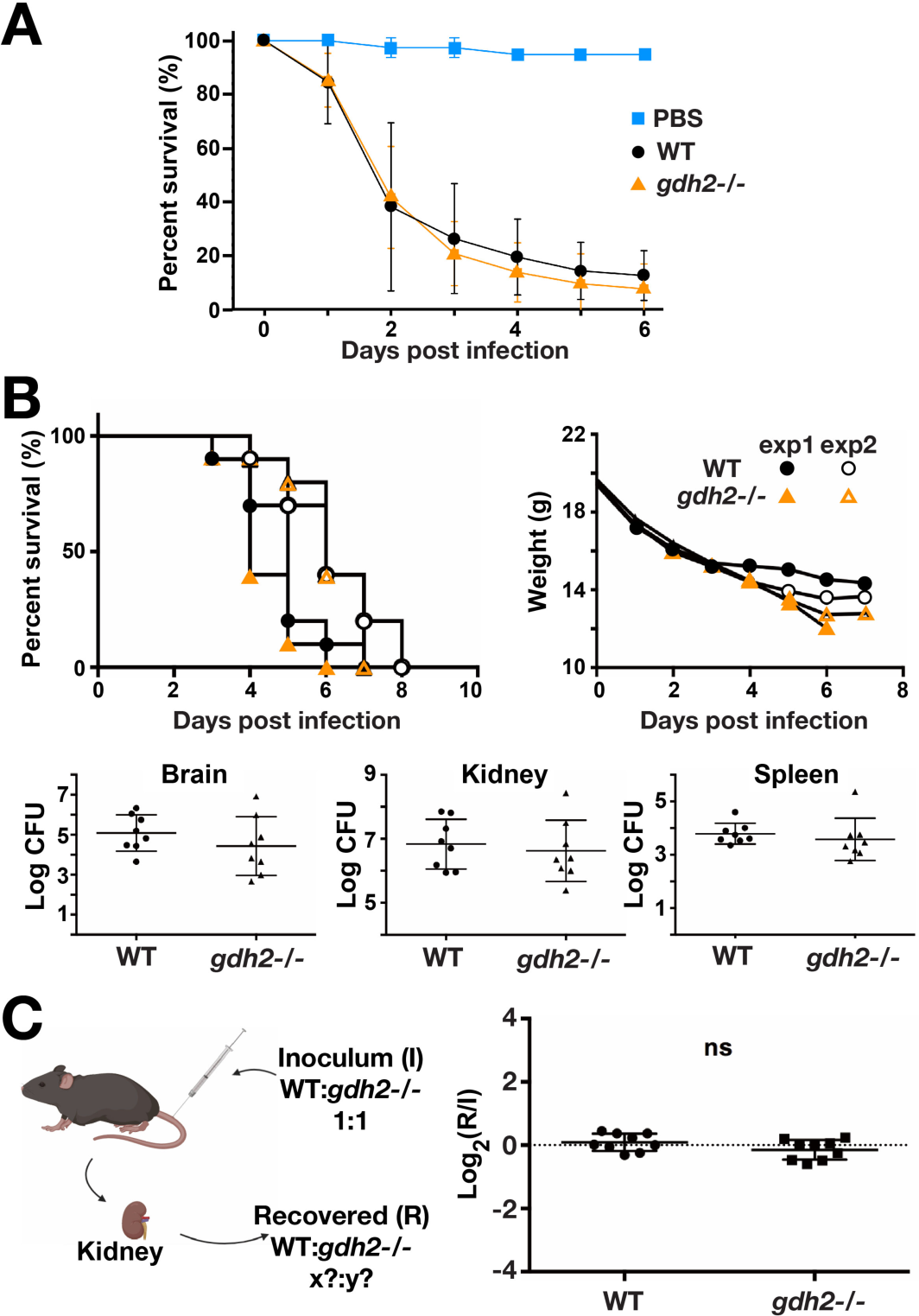
Virulence of wildtype and *gdh2-/- C. albicans* in *Drosophila* and murine systemic infection models. (A) *D. melanogaster Bom*^Δ*55C*^ flies were infected with wildtype (SC5314) or *gdh2-/-* (CFG279) cells as indicated, and the survival of flies was followed for six days. Each curve represents the average of a minimum of three independent infection experiments (20 flies/strain) performed on different days. (B) Groups of C57BL/6 mice (n=10) were infected via the lateral tail vein with 3x10^5^ CFU of *C. albicans* wildtype or *gdh2*-/- cells (upper panels) and survival (left) and weight loss (right) was monitored at the timepoints indicated. Survival curves from two independent experiments were statistically analyzed by the Kaplan-Meier method (a log-rank test, GraphPad Prism), no significant difference. The fungal burden (lower panels) in brain (left), kidney (middle), and spleen (right) extracted from mice 3 days post infection. Each symbol represents a sample from an individual mouse and results were compared by Student *t*-test, no significant difference. (C) Competition assay; mice were infected via the tail vein with an inoculum (I) comprised of an equal number of wildtype (SC5314) and *gdh2-/-* (CFG279), 1:1. At 3 days post infection, the abundance and genotype of fungal cells recovered from kidneys was quantitated and the ratio of wildtype:*gdh2-/-* recovered (R) was determined. The significance of the log_2_(R/I) values was assessed using an unpaired *t*-test, no significant difference (ns).

To further assess the importance of Gdh2 in virulence in a more complex host, we used a tail vein infection model using C57BL/6 mice. Two groups of mice (n = 10) were challenged with 3 x 10^5^ wildtype or *gdh2*-/- cells and survival was monitored for a period of up to 8 days. Similar to the fly model, we observed that the loss of Gdh2 activity did not attenuate virulence (**Fig. 7B**); the *gdh2*-/- mutant exhibited survival indistinguishable from wildtype. Consistently, the fungal burden of *gdh2*-/ cells in the brain, kidney and spleen of infected mice 3 days post-infection did not significantly differ to mice infected with wildtype (**Fig. 7C**). Next we performed a competition assay; equal numbers of wildtype and *gdh2*-/- cells were intravenously injected in mice and the ratio (R) of wildtype to *gdh2*-/- cells recovered from kidneys 3 days post-infection was determined. Consistent to our findings in mice individually infected with each strain, the ratio of recovered cells did not significantly differ to that of the inoculum ratio (I) (**Fig. 7C**). Together, our results indicate that Gdh2 is not required for virulence, and that the loss of Gdh2 activity does not create a selective disadvantage or significantly impair growth in infected model host systems.

## DISCUSSION

In this work, we identified a major metabolic step that endows *C. albicans* with the capacity to increase the extracellular pH by ammonia extrusion in the presence of amino acids. We have shown that under *in vitro* growth conditions, environmental alkalization is dependent upon *GDH2*, a gene that encodes the mitochondrial-localized glutamate dehydrogenase. Strikingly, the data clearly show that despite its unambiguous role in alkalization *in vitro*, *GDH2* remained dispensable for the induction of hyphal growth and escape of *C. albicans* from macrophages, and also dispensable for virulence in intact hosts. Our results, consistent with a recent report (25), suggest that phagosomal alkalization is unlikely to be a defining event required for hyphal initiation of *C. albicans* in phagosomes of engulfing macrophages.

Despite its role as a key enzyme of central nitrogen metabolism, the *gdh2-/-* mutant exhibited only a modest growth defect on synthetic glucose or glycerol media containing, glutamate as sole nitrogen source (**Fig. S1C**). Consistent with proline being catabolized to glutamate in a linear pathway mediated by Put1 and Put2, a similar modest growth defect of *gdh2-/-* was observed when proline was the sole source of nitrogen (**Fig. S1C**). However, when glucose or glycerol is removed from the media (i.e., YNB+CAA), and amino acids serve as both carbon and nitrogen sources, the *gdh2*-/- mutant demonstrated a striking growth defect (**Fig. 1B, 1C, S2A**). Under these conditions, the impaired growth of cells lacking Gdh2 is likely the consequence of diminished levels of α-ketoglutarate. In the absence of Gdh2, cells must rely entirely on the TCA cycle to generate α-ketoglutarate required to support *de novo* biosynthetic needs, e.g., amino acid biosynthesis. In the absence of glucose or glycerol, the supply of acetyl-CoA derived from pyruvate becomes limiting, stalling the TCA cycle, and consequently, the growth of *gdh2*-/- cells.

Similar to *S. cerevisiae*, the Gdh2-catalyzed reaction is the primary source of extruded ammonia in *C. albicans*. This conclusion is based on the following key observations: 1) a mutant strain lacking *GDH2* (*gdh2*-/-) is unable to alkalinize media with amino acids as carbon and nitrogen source (i.e., YNB+CAA) and under non-repressing glucose conditions (**Fig. 1C**); 2) ammonia extrusion is impaired in *gdh2*-/- mutant (**Fig. 1D**); and 3) the Gdh2-catalyzed reaction responsible for alkalization occurs in the mitochondria, and is subject to glucose repression and inhibited by inhibitors of mitochondrial respiration (**Fig. 2**).

The capacity of glucose to repress mitochondrial activity (15) and Gdh2 expression (**Fig. 3C**) may explain why alkalization is only observed when glucose becomes limited or replaced by glycerol. The induced levels of Gdh2 expression observed upon phagocytosis is consistent with the apparent low levels of glucose in phagosomes (27). Together with an expected surge in glutamate coming from the catabolism of arginine or proline (15); Put2 levels are upregulated in phagocytized *C. albicans*, which requires proline binding to the transcription factor Put3 (15, 28). In the presence of arginine as sole nitrogen and carbon source (i.e., YNB+Arg), the role of proline catabolism in alkalization is essential since it is the primary route to generate glutamate in the mitochondria. However, when other amino acids are present (i.e., YNB+CAA) such as glutamine, alanine, and aspartate (29), these amino acids can be converted directly to glutamate bypassing the requirement for the proline catabolic pathway. The fact that the *put1-/-* strain still showed a weaker alkalization compared to wildtype in YNB+CAA indicates that in the absence of glucose, proline functions a preferred carbon source, which is likely due its catabolism is efficiently coupled to the generation of ATP.

An unanswered question is how ammonia generated in the mitochondria is extruded to the external environment. It is possible that ammonia (NH_3_), known to be membrane permeable, diffuses across the inner mitochondrial membrane, moving towards the more acidic inner membrane space where it likely becomes protonated to ammonium. Ammonium then moves to the cytosol. Although the dissociation of ammonium to ammonia is not favored at the pH of the cytosol (pH ∼7), the small amount of ammonia that forms can rapidly diffuse across the PM out of cells as long as the external environment is acidic. Hence, the ability of the ammonia generated by Gdh2 will likely be the consequence of Pma1 activity, the major proton pumping ATPase in the PM. Alternatively, and according to several reports, putative ammonia transport proteins, the Ato family of plasma membrane proteins, are thought to facilitate ammonia export in *C. albicans* (11). Supporting this notion, the deletion of *ATO5* significantly delays alkalization. Interestingly, the requirement for the Ato proteins suggests that the species traversing the plasma membrane from within the cell is either charged or polar, thus, it is likely that the transported species is ammonium (NH4^+^) and coupled to H import as previously suggested in yeast (30, 31). Since cytoplasmic pH is tightly regulated, the conundrum persists as to how extruding ammonium can facilitate steady-state alkalization. The underlying mechanism of how Ato proteins facilitate alkalization needs to be precisely defined and placed in context to the fact that ammonia can readily diffuse through membranes, and in so doing, is expected to move directionally towards acidic environments.

Due to its central role in nitrogen metabolism, it was surprising that the inactivation of *GDH2* did not affect the capacity of *C. albicans* to form hyphae in the phagosome of macrophage, which is thought to contain amino acids as primary energy sources. This is opposite to what we observed in strains lacking *PUT1* and/or *PUT2,* which show phagosome-specific defect in hyphal growth (15). On amino acid-rich Spider and Lee’s medium, containing 1% mannitol and 1.25% glucose as primary carbon sources, respectively, the *gdh2*-/- mutant also did not show a filamentation defect. On Spider, but not on Lee’s media, both *put1-/-* and *put2-/-* mutants have noticeable defects on formation of invasive filaments despite forming wrinkled colonies (our unpublished data). Together, these observations suggest that when glucose is limiting, the energy obtained by the catabolism of proline to glutamate suffices to induce and support hyphal growth; the additional energy derived from the NADH generated by the Gdh2-catalyzed deamination of glutamate is not required. The recent work by Westman et al., demonstrating that phagosomal alkalization is an effect of hyphal expansion and not the underlying trigger that causes filamentation (25), aligns well with our observations. Accordingly, Westman et al. proposed that the step-wise alkalization of the phagosome could be attributed to proton leakage out of the compartment due to the transient physical stress imposed by hyphal expansion. Also, hyphal formation was found to start prior to a measurable change in pH (pH ∼ 5-6), suggesting that alkalization is not the primary stimulus triggering hyphal formation in the phagosome. Beyond the realm of the phagosome, our work also suggests that environmental alkalization via ammonia extrusion, a mechanism that is thought to facilitate virulence of fungal pathogens (32), is dispensable for pathogenesis of *C. albicans* (**Fig. 7**) requiring us to rethink the specific role of alkalization in fungal virulence.

We confirmed that *DUR1,2* does not significantly contribute to alkalization. *DUR1,2* is under tight regulatory control by NCR, and thus is not expressed under growth conditions in the presence of preferred amino acids. Our results are inconsistent with previous suggestions that Dur1,2 activity significantly contributes to alkalization (12). The usual growth media (i.e., YPD or SD) used for *C. albicans* propagation and the standard mammalian cell culture medium (i.e., DMEM or RPMI) used for co-culturing fungal cells with macrophages are all very rich in amino acids, certainly conditions that repress *DUR1,2* expression (33). In co-culture experiments, we observed that hyphal initiation (i.e., germ tube formation) following phagocytosis is very rapid, occurring as early as 15-20 min following phagocytosis even in experiments that were carried out in a neutral buffer (i.e., HBSS) suggesting signaling cascades driving this switch are activated at a much earlier time.

Finally, we note that time-lapse microscopy has a distinct advantage over endpoint microscopy of fixed co-cultures, since it enables the spatio-temporal dynamics of hyphal formation of both wildtype and *gdh2-/-* mutant to be accurately followed inside the same macrophage. Time-lapse microscopy allowed us to observe that non-phagocytized fungal cells that remained external even after excessive washing can filament resulting in the false impression that the fungal cells are escaping from the macrophage.

Counteracting the antimicrobial assault in the macrophage phagosome is crucial for *C. albicans* survival and dissemination. Due to acidic phagosomal microenvironment, phagosomal alkalization via ammonia extrusion is not surprising at all. However, given that *C. albicans* is retained in a hostile microenvironment essentially devoid of key nutrients required for growth, it is of paramount importance for *C. albicans* to be able to synthesize the cellular components required to counteract these stresses. For example, the highly polarized hyphal growth requires a lot of ATP to drive actin polymerization and also the proton extrusion process mediated by Pma1 to regulate intracellular pH when extracellular pH is low requires ATP. Thus, on a bioenergetic standpoint, and considering the energy-demanding nature of hyphal function, the alkalization process does not seem adequate for an explanation as to how *C. albicans* is able to support hyphal formation in a hostile microenvironment with restricted nutrient content. The ability to undergo dimorphic transitions to hyphal growth and the distention of phagosomal membrane apparently has major physiological consequence that affects the capacity of macrophage killing. The findings documented here illuminate and further the understanding of a major feature of the innate immune surveillance arsenal required for integrity of the human host, however, more work is clearly needed to understand the spatio- temporal dynamics of hyphal formation of *C. albicans* in the macrophage phagosome.

## METHODOLOGY

### Organisms, culture media, and chemicals

Strains listed in Table S1 were routinely cultivated in YPD agar medium (1% yeast extract, 2% peptone, 2% glucose, 2% Bacto agar) at 30 °C after recovery from -80 °C glycerol stock. Where needed, YPD medium was supplemented with 25, 100 or 200 µg/ml nourseothricin (Nou; Jena Biosciences, Jena, Germany). Also, where indicated, the glucose in YPD is lowered to 0.2% (YPD_0.2%_) or replaced with 1% glycerol (YPG) or 2% mannitol (YPM). Specific growth assays were carried out in synthetic minimal medium containing 0.17% yeast nitrogen base without ammonium sulfate and amino acids (YNB; Difco), supplemented with the indicated amino acid (10 mM) as sole nitrogen source and the indicated carbon source, and buffered (pH = 6.0, 50 mM MES).

### Alkalization assays

Alkalization was assessed using YNB+CAA medium (0.17% YNB, 1% of casamino acids (CAA; Sigma) containing 0.01% Bromocresol Purple (BCP; Sigma) as pH indicator; the pH was set at 4.0 using 1 M HCl. Where indicated, YNB+Arg was used, which contains 10 mM arginine instead of 1% CAA as sole nitrogen and carbon source. Also, where indicated, YNB+CAA medium was supplemented with glycerol (1%) or glucose (2% or 0.2%). Cells from overnight YPD cultures were harvested, washed at least twice in ddH_2_O, and then suspended at an OD_600_ ≈ 0.05 unless otherwise indicated. Cultures were grown with vigorous agitation at 37 °C. Where appropriate Antimycin A or Chloramphenicol was added at the concentrations indicated. Assays on solid media (2% agar), 5 μl aliquots of washed cell suspensions (OD_600_ ≈ 1) were spotted onto the surface of media in a 6- well microplate and then grown at 37 °C for up to 72 h.

### CRISPR/Cas9 Mediated Gene Inactivation

CRISPR/Cas9 was used to simultaneously inactivate both alleles of *GDH2* (C5_02600W) (34, 35). Synthetic guide RNAs (sgRNAs), repair templates (RT), and verification primers used for gene editing are listed in Table S2. Briefly, 20-bp sgRNAs primers (p1/p2), designed according to (36), were ligated to *Esp*3I (*Bsm*BI)-restricted and dephosphorylated pV1524 creating pFS108. The CRISPR/Cas9 cassette (100 ng/μl) of pFS108, with sgRNA targeting *GDH2*, was released by *Kpn*I and *Sac*I restriction and introduced into *C. albicans* together with a PCR-amplified RT (p3/p4; 100 ng/μl) containing multiple stop codons and a diagnostic *Xho*I restriction site (plasmid:repair template volume ratio of 1:3). *C. albicans* transformation was performed using the hybrid lithium acetate/DTT-electroporation method by Reuss, et al. (37). After applying the 1.5 kV electric pulse, cells were immediately recovered in YPD medium supplemented with 1 M sorbitol for at least 4 hours, and then plated on YPD-Nou plates (200 µg/ml). Nou-resistant (Nou^R^) transformants were re-streaked on YPD- Nou plates (100 µg/ml) and screened for the ability to alkalinize YNB+Arg media. DNA was isolated from transformants exhibiting an alkalization defect and subjected to PCR-restriction digest (PCR-RD) verification using primers (p5/p6) and *Xho*I restriction enzyme. The *gdh2-/-* clones (CFG277 and CFG278) were grown overnight in YPM to pop-out the CRISPR/Cas9 cassette. Nou sensitive (Nou^S^) cells were identified by plating on YPD supplemented with 25 µg/ml Nou (37), resulting in strains CFG279 and CFG281.

### Reporter Strains

For C-terminal GFP tagging of Gdh2, an approximately 2.8 kB of PCR cassette was amplified from plasmid pFA-GFPγ-*URA3* (38) using primers (p7/p8). The amplicon was purified and then introduced into CAI4 (*ura3*/*ura3*). Transformants were selected on synthetic complete dextrose (CSD) plate lacking uridine. The correct integration of the GFP reporter was assessed using PCR (p5/p9), immunoblotting, and fluorescence microscopy. CFG275, a *gdh2*-/- strain constitutively expressing RFP, was constructed by introducing a *Kpn*I/*Sac*I fragment from pJA21 containing the P*_ADH1_-*RFP-Nou construct (39) into CFG279; transformants were selected on YPD agar with 200 µg/ml Nou. RFP-positive clones were verified by PCR (p11/p12) and fluorescence microscopy.

### Ammonia release assay

Quantification of volatile ammonia release was performed in accordance to the modified acid trap method by Morales et al. (40). Briefly, a 2 μl aliquot of OD_600_ ≈ 1 cell suspension was spotted onto each well of a 96-well microplate containing 150 µl of YNB+CAA solid medium supplemented with 0.2% glucose and buffered to pH = 7.4 with 50 mM MOPS. The spotted microplate was inverted and then precisely positioned on top of another microplate in which each well contains 100 µl of 10 % (w/v) citric acid. Plates were sealed by parafilm and then incubated at 37°C for 72 h after which, the citric acid solution was sampled for ammonia analysis using Nessler’s reagent (Sigma-Aldrich). The solution was diluted 10-fold in 10% citric acid and then a 20-µl aliquot was added to 80 µl Nessler’s reagent on another microplate. After a 30 min incubation period at room temperature, OD_400_ was measured using Enspire microplate reader. The level of ammonia entrapped in the citric acid solution was calculated based on ammonium chloride (NH_4_Cl) standard curve.

### Immunoblotting

For Gdh2 level analysis, cells expressing Gdh2-GFP (CFG273) were grown in liquid YPD for overnight at 30°C and then washed 3X with ddH_2_O. Cells were diluted in the indicated alkalization media at OD_600_ ≈ 2 and then incubated continuously in a rotating drum for 6 h at 37°C with sampling performed every 2 h. In each sampling point, cells were harvested, washed once with ice-cold ddH_2_O, and then adjusted to OD_600_ ≈ 2. Whole cell lysates were prepared using sodium hydroxide/ trichloroacetic acid (NaOH/TCA) method as described previously with minor modifications (41). Briefly, 500 µl of adjusted cell suspension were added to tube containing 280 µl of ice-cold 2 M NaOH with 7% ß-Mercaptoethanol (ß-Me) for 15 min. Proteins were then precipitated overnight at 4°C by adding the same volume of cold 50% TCA. Protein pellets were collected by high-speed centrifugation at 13,000 rpm for 10 min (4°C) and then the NaOH/TCA solution completely removed. The pellets were resuspended in 50 µl of 2X SDS sample buffer with additional 5 µl of 1 M Tris Base (pH = 11) to neutralize the excess TCA. Samples were denatured at 95-100°C for 5 min before resolving the proteins in sodium dodecyl sulfate-polyacrylamide gel electrophoresis (SDS-PAGE) using 4- 12% pre-cast gels (Invitrogen). Proteins were analyzed by immunoblotting on nitrocellulose membrane according to standard procedure. After transfer, membranes were blocked using 10% skimmed milk in TBST (TBS + 0.1% Tween) for 1 h at room temperature. For Gdh2-GFP detection, membranes were first incubated with mouse anti-GFP primary antibody at 1:2,000 dilution (JL8, Takara) for overnight at 4°C. For the detection of the primary antibody, an HRP-conjugated goat anti-mouse secondary antibody (Pierce) was used. For loading control, α-tubulin was detected with rat monoclonal antibody conjugated to HRP [YOL1/34] (Abcam). For both secondary antibody and loading control, antibodies were used at 1:10,000 dilution in 5% skimmed milk in TBST incubated for 1 h at room temperature. Immunoreactive bands were visualized by enhanced chemiluminescent detection system (SuperSignal Dura West Extended Duration Substrate; Pierce) using ChemiDoc MP system (BioRad).

### Filamentation assay

Filamentation in solid Spider (42) or Lee’s (43) media was performed as described (44). Cells from overnight YPD liquid cultures were harvested, washed 3X with sterile PBS, adjusted to OD_600_ ≈ 1 and then 5 µl of cell suspensions were spotted onto the indicated media. Plates were allowed to dry at room temperature before incubating at 37 °C as indicated.

### Macrophage culture

RAW264.7 murine macrophage cells (ATCC TIB-71) and primary bone marrow-derived macrophages (BMDM) were cultured and passaged in complete RPMI medium supplemented with 10% fetal bovine serum (FBS), 100 U/ml penicillin and 100 mg/ml streptomycin (referred to as R10 medium in the text) in a humidified chamber set at 37°C with 5% CO_2_. For BMDM differentiation, bone marrows collected from mouse femurs of C57BL/6 wildtype mice (7- to 9- week old) were mechanically homogenized and resuspended in R10 medium supplemented with 20% L929 conditioned media (LCM). Differentiation was carried out initially for 3 days before boosting the cells with another dose of 20% LCM until harvested.

BMDM were used 7-10 days after differentiation.

### *C. albicans* killing assay

To assess candidacidal activity by BMDM, we co- cultured *C. albicans* wildtype and *gdh2*-/- mutant with BMDM and then assessed colony forming units (CFU) following co-incubation. About 16-24 h prior to co- culture, differentiated BMDM were collected by scraping, counted, and then seeded at 1 x 10^6^ cells/well into a 24-well microplate. *C. albicans* cells from overnight YPD cultures were collected by centrifugation, washed 3X with sterile PBS, and then added to macrophages at MOI 3:1 (C:M). The plates were briefly centrifuged at 500 x g for 5 min to collect the fungal cells at the bottom of each well and then co- cultured for 2 h in a humidified chamber. After co-culture, each well was treated with 0.1% Triton X-100 for 2 min followed by vigorous pipetting to lyse the macrophage and release the fungal cells. Each well was rinsed seven times (7X) with ice-cold ddH_2_O and collected in a 15-ml conical tube. Lysates were serially diluted and then plated onto YPD. Plates containing colonies between 30-300 were counted. The candidacidal activity (% killing) of BMDM was defined as [1 - (CFU with macrophage/CFU of initial fungal inoculum)] x 100 (45).

### Confocal microscopy

For subcellular localization of Gdh2, cells expressing Gdh2- GFP (CFG273) from log-phase YPD cultures were harvested, washed 3X with ddH_2_O, and then grown in SED_0.2%_ (10 mM glutamate and 0.2% glucose) and YNB+CAA for 24 h at 37°C at a starting OD_600_ ≈ 0.05. Cells from each culture were harvested, washed once with PBS, and then stained with 200 nM (in PBS) of the mitochondrial marker, MitoTracker Red (MTR; Molecular Probes) for 30 min at 37°C. After staining, the cells were collected again and resuspended in PBS before viewing the cells using confocal microscope (LSM800, 63x oil) in the green and red channels.

For the investigation of hyphal formation in the phagosome, we co-cultured individual strains with RAW264.7 macrophages. For easier visualization, we pre- stained the fungal cells with fluorescein isothiocyanate (FITC). Briefly, fungal cells were harvested from YPD overnight cultures, washed once with PBS, and then adjusted to OD_600_ ≈ 10. Cells from 1 ml of adjusted cell suspension were collected and then incubated with FITC solution (1 mg/ml in 0.1 M NaHCO_3_ buffer, pH = 9.0) for 15 min at 30 °C before extensive washing in PBS (3X). Macrophages were seeded at 1 x 10^6^ cells into a 35 mm glass bottom imaging dish (ibidi) and were allowed to adhere overnight (16-24 h). FITC stained fungal cells were added at a MOI of 1:1 (C:M), and after 20 min, the co-cultures were washed extensively with HBSS (5X) to remove non-phagocytosed fungal cells before adding CO_2_- independent medium (Gibco). Co-cultures were further incubated for 1.5 h in temperature-controlled chamber (37 °C) of the LSM800 confocal microscope prior to imaging cells (63x objective).

### Time lapse microscopy (TLM)

Unless otherwise indicated, all TLM experiments performed in this paper used the Zeiss Cell Observer inverted microscope equipped with temperature control chamber (37 °C) and appropriate filters to detect fluorophores. For the analysis of Gdh2 expression during alkalization on solid media, cells expressing Gdh2-GFP (CFG273) were collected from overnight YPD culture, washed 3X with ddH_2_O, and then adjusted to a cell density of OD_600_ ≈ 0.05. To make a YNB+CAA agar slab on which to grow the cells, a 100 µl molten YNB+CAA agar was placed on top of a flame-sterilized slide and then spread evenly to make a thin agar film. The agar was allowed to congeal at room temperature before spotting a 2 µl aliquot of adjusted cell suspension and then covered with a coverslip. Single cells were located and then the GFP expression was followed every hour in the green (GFP) channel alongside DIC for 6 h.

For Gdh2-GFP expression during macrophage interaction, the same strain (CFG273) was co-cultured with either RAW264.7 or BMDM macrophage pre-stained with LysoTracker Red DND-99 (LST; Thermo Scientific) that marks the acidic compartments inside the macrophage. Macrophages were seeded at 1 x 10^6^ cells into a 35 mm glass bottom imaging dish and were allowed to adhere overnight (16- 24 h). Prior to co-culture, medium was removed and then replaced with CO_2_- independent medium containing 200 nM of LST. Macrophages were stained for at least 30 min at 37°C. For fungal cell preparation, cells from overnight YPD cultures were harvested, washed 3X with sterile PBS, and then added to macrophages at MOI of 1:1 (C:M). Interaction was followed every 1.5 min for ∼3 h (with RAW cells) and 2 min for ∼4 h (with BMDM) in the DIC, green (Gdh2-GFP) and red (LST) channels. Movies were saved at 10 fps.

For competition assay in the same macrophage (BMDM) co-culture system, wildtype cells constitutively expressing GFP (SCADH1G4A) and *gdh2*-/- strain expressing RFP (CFG275) from overnight YPD cultures were collected by centrifugation, washed 3X with sterile PBS, and then diluted to OD_600_ ≈ 1. Cells were mixed 1:1 (v/v) in a sterile Eppendorf tube and then vortexed. Prior to co-culture, BMDM (1 x 10^6^) seeded on imaging dish were washed 2X with HBSS to remove the growth medium. A 100 μl aliquot of mixed fungal cells (∼3 x 10^6^ cells) were added to the dish (MOI of 3:1, C:M) and phagocytosis was carried out in HBSS for approximately 30 min in the humidified chamber. Co-cultures were washed at least 5X with HBSS and 1X with CO_2_-independent medium to remove non-phagocytosed fungal cells. CO_2_-independent medium was added to the dish and TLM was carried out at 37°C in the DIC, green (wildtype) and red (*gdh2*-/-) channels. Images were acquired every 2 min for ∼5 h and then saved as movie at 10 fps.

### Murine systemic infection model

Groups of female C57BL/6 mice, aged 6–8 weeks, were purchased from Beijing Vital River Laboratory Animal Technology Co., Ltd. These mice were housed in individual ventilated cages in a pathogen-free animal facility at Institut Pasteur of Shanghai, Chinese Academy of Sciences. In each of the mouse studies, the animals were assigned to the different experimental groups. Infections were performed under SPF conditions. All animal experiments were carried out in strict accordance with the Regulations for the Administration of Affairs Concerning Experimental Animals issued by the Ministry of Science and Technology of the People’s Republic of China, and approved by IACUC at the Institut Pasteur of Shanghai, Chinese Academy of Science with an approval number P2018050. Wildtype SC5314 and *gdh2-/-* mutant *C. albicans* cells were inoculated into YPD broth and grown overnight at 30 °C. Cells were harvested and washed three times with phosphate-buffered saline (PBS), and counted using hemocytometer. For each strain, mice (n=10) were injected via the lateral tail vein with 3x10^5^ CFU of *C. albicans* cells. The mice were monitored once daily for weight loss, disease severity and survival. The fungal burden was assessed by counting CFU. The survival curves were statistically analyzed by the Kaplan-Meier method (a log-rank test, GraphPad Prism). Competitive bloodstream infections were performed using equal numbers of SC5314 and *gdh2-/-*mutant cells, i.e., with an inoculum (I) ratio of 1:1. At 3 days post infection the abundance and genotype of fungal cells recovered from kidneys was determined and the ratio of wildtype:*gdh2-/-* in kidneys was calculated (R). Cells lacking *gdh2-/-* cannot grow on selective YNB+Arg medium. The log_2_(R/I) values was compared using unpaired t-test.

## Author contribution statements

F.G.S.S. and P.O.L. conceived and designed the experiments. F.G.S.S., K.R., T.J., M.W., and N.H. performed experiments. F.G.S.S., C.C., and N.N.L. and P.O.L. supervised the experimental work. F.G.S.S., K.R., M.W., N.H., C.C., T.J., N.N.L., and P.O.L. analyzed the data and prepared the figures. F.G.S.S. and P.O.L. wrote the paper. P.O.L. acquired the main funding and all authors critically reviewed and approved the manuscript.

## Acknowledgments

The authors would like to thank the members of the Claes Andréasson, Sabrina Büttner, Roger Karlsson and Per Ljungdahl laboratories (SU) for their constructive comments throughout the course of this work. Gratitude is extended to Valmik Vyas and Gerard Fink (MIT, Cambridge, MA, USA) for providing the CRISPR/Cas9 cassettes and Joachim Morschhäuser (Universität Würzburg, Germany) for supplying strains and for fruitful discussions. We also thank Stina Höglund, the Imaging Facility-Stockholm University (IFSU), for assistance in microscopy. We would also like to acknowledge Andreas Ring (SU), Joachim Morschhäuser (Universität Würzburg) and Johannes Westman (Hospital for Sick Children, Toronto, ON, CA) for constructive comments of the manuscript. This work was supported by EU grant MC-ITN-606786 (ImResFun) and grants from the Swedish Research Council VR-2015-04202 and 2019-01547 (POL).

## Supporting Information

### Supporting Information – Figures

**Fig. S1.**
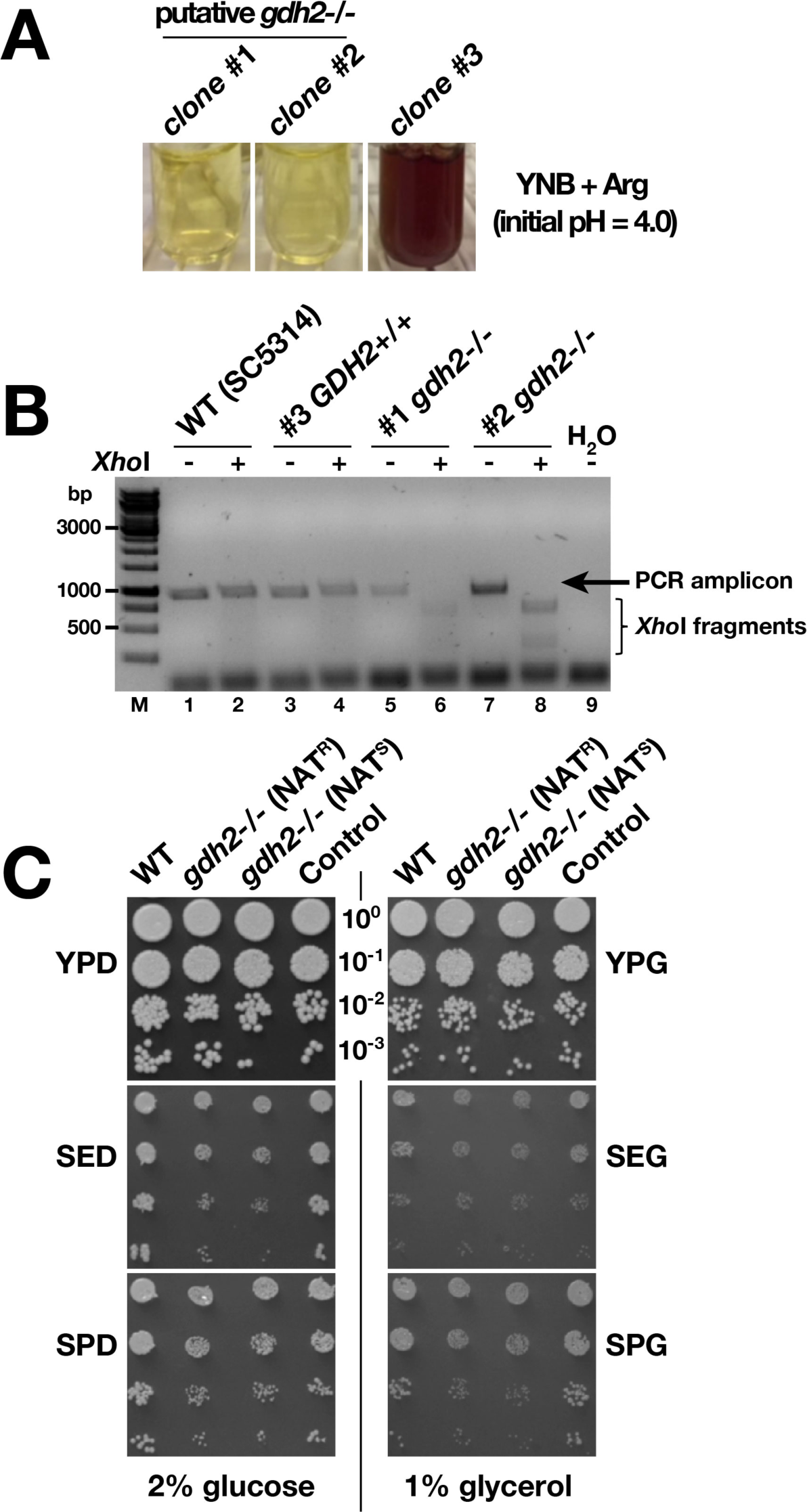
CRISPR/Cas9-mediated gene inactivation of *GDH2* in *C. albicans*. (A) A purified *Kpn*I/*Sac*I fragment from pFS108, harboring *GDH2*-specific sgRNA, and PCR generated repair template (RT) were introduced into wildtype strain SC5314 by electroporation. Nou^R^ transformants were pre- screened in YNB+Arg medium containing the pH indicator bromocresol purple; the initial pH was 4.0. Three Nou^R^ colonies were picked for further analysis. Clones #1 and #2 grew poorly and were unable to alkalinize the media; clone #3 grew and alkalinized the media. (B) Genomic DNA, isolated from the three clones, was used as template for PCR amplification of the targeted *GDH2* locus; ddH_2_O was used as negative control. Restriction of the amplified ≈900 bp fragment by *Xho*I is diagnostic for successful mutagenesis. Strains, clone #1 (CFG277) and clone #2 (CFG278) carry inactivated *gdh2-/-* alleles. (C) *GDH2* is not essential but required for robust growth on glutamate or proline as sole nitrogen source. Five microliters of serially diluted wildtype (SC5314), *gdh2-/-* NAT^R^(CFG277), *gdh2-/-* NAT^S^(CFG279), and control (CFG182) cells were spotted on yeast peptone (YP), synthetic glutamate (SE) and synthetic proline (SP) media containing either 2% glucose (D) or 1% glycerol (G) as carbon source. The plates were incubated for 48 h at 30°C and photographed.

**Fig. S2.**
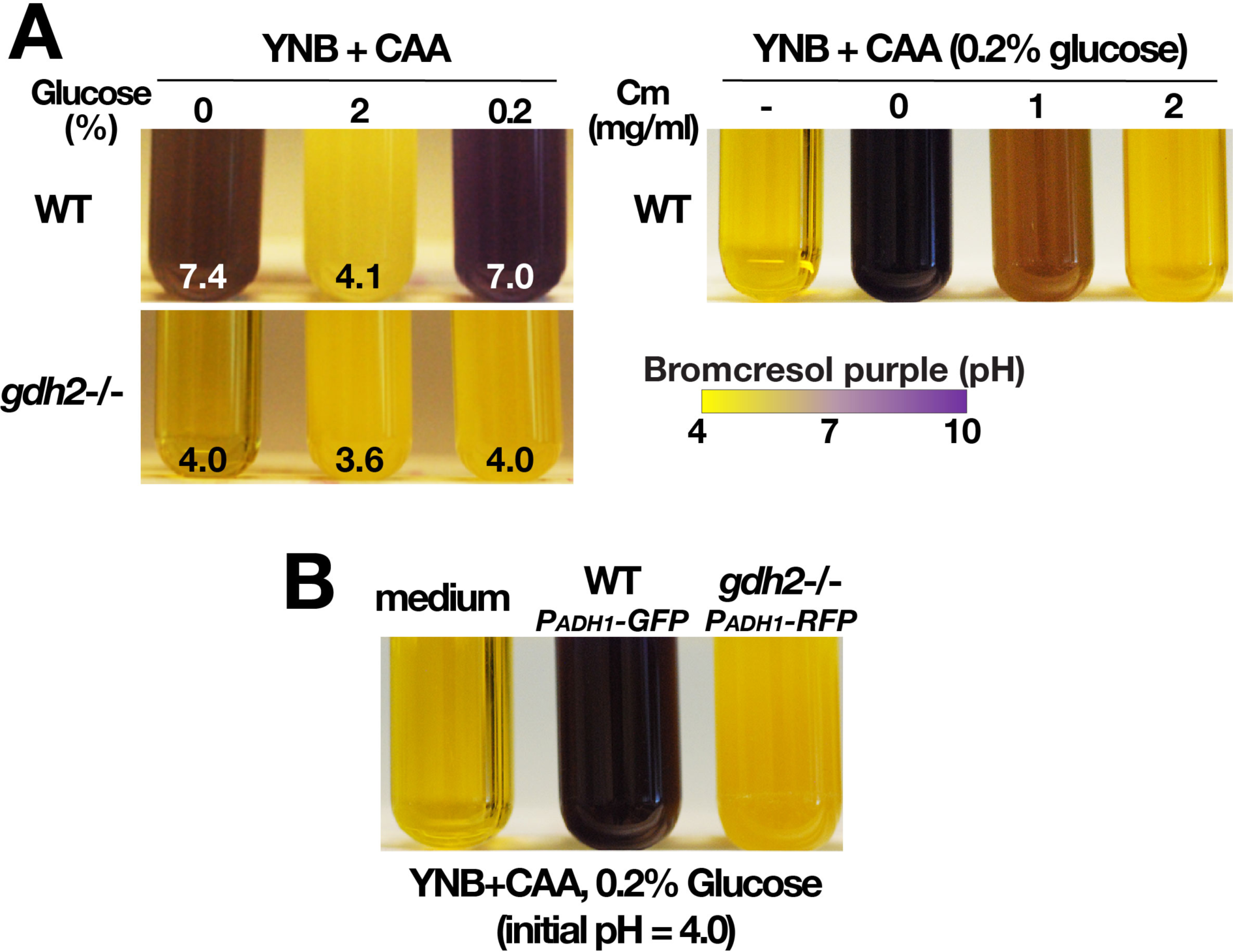
Growth characteristics of wildtype and *gdh2-/-* strains in liquid YNB+CAA with and without glucose and chloramphenicol (Cm) (A) Gdh2-dependent alkalization is sensitive to glucose (Left panel). YPD grown wildtype (WT, SC5314) and *gdh2-/-* (CFG279) cells were collected, washed, and diluted to an OD_600_ of 0.05 in YNB+CAA with 0, 2 or 0.2% glucose as indicated. The cultures were grown under vigorous agitation at 37 °C for 16 h and the pH was measured (the initial pH was 4.0; the values indicated are the average of three replicate cultures). Alkalization is linked to mitochondrial function (Right panel). Wildtype cells (SC5314) from overnight YPD cultures were washed and diluted to OD_600_ ≈ 0.1 in liquid YNB+CAA (0.2% glucose) with the indicated concentrations of mitochondrial translation inhibitor chloramphenicol. Cultures were grown at 37 °C under vigorous agitation for 16 h. (B) Phenotypic validation of the reporter strains used in macrophage co-cultures. Growth of wildtype (WT; *P_ADH1_-GFP*; SCADH1G4A) and *gdh2*-/- (*P_ADH1_-RFP*, CFG275) cells in YNB+CAA supplemented with 0.2% glucose. Cultures were grown for 16 h at 37 °C.

### Supporting Information - Tables

**Table S1.**
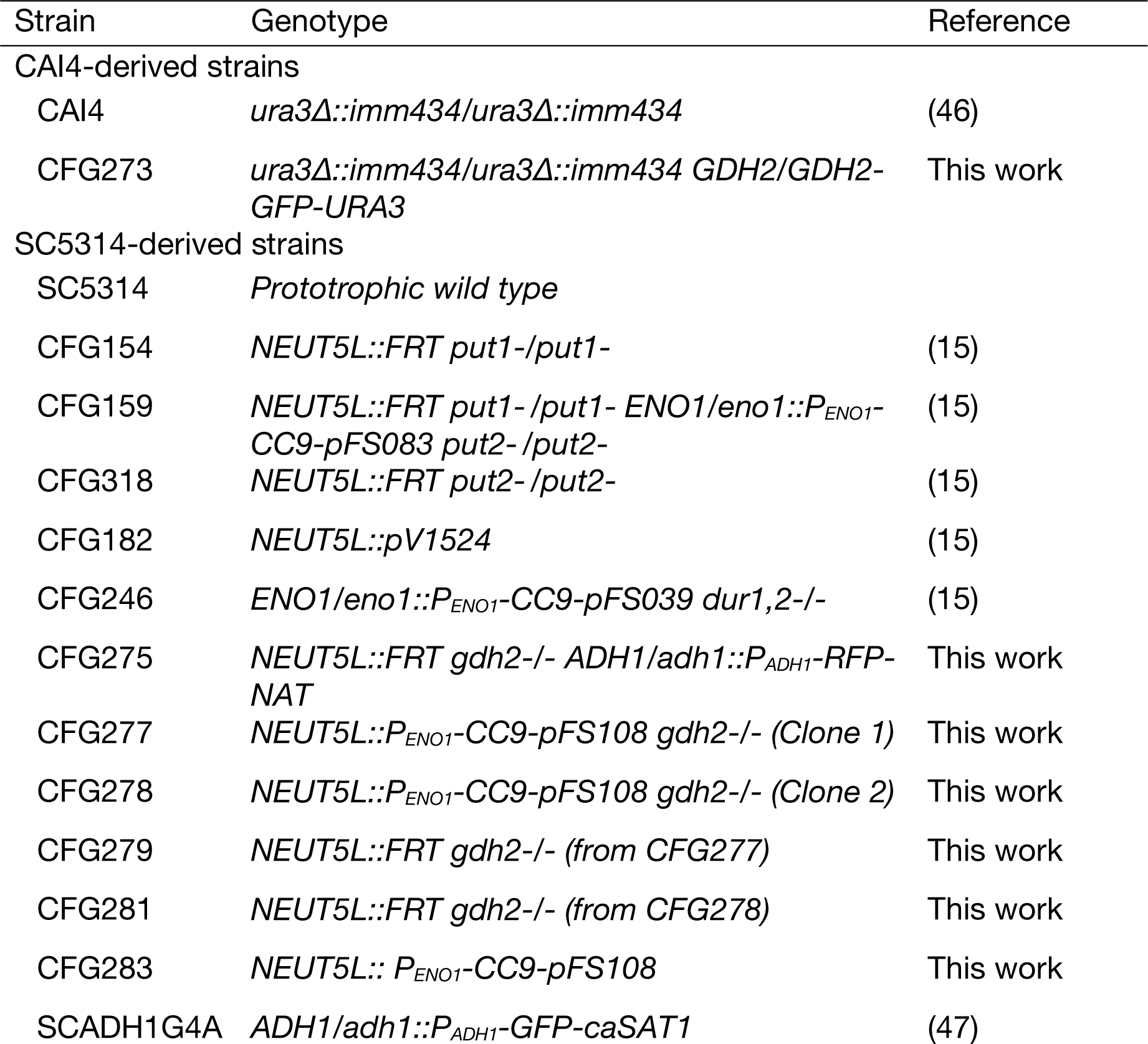
Strains used in this study.

**Table S2.**
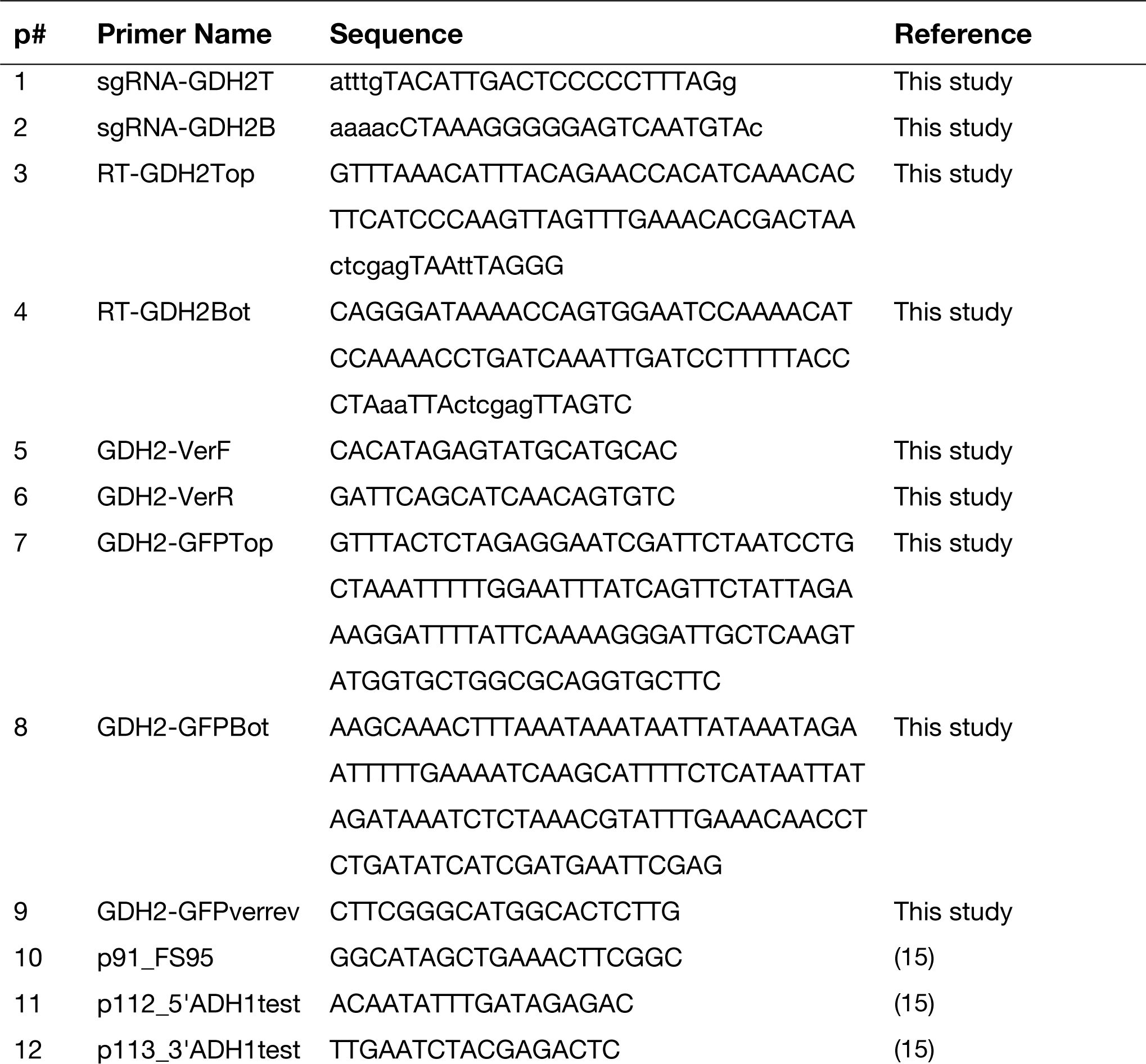
Primers used in this study

## REFERENCES

1. Kullberg BJ, Arendrup MC. Invasive Candidiasis. N Engl J Med. 2016;374(8):794–5.

2. Moyes DL, Naglik JR. Mucosal immunity and *Candida albicans* infection. Clin Dev Immunol. 2011;2011:346307.

3. Berman J. Morphogenesis and cell cycle progression in *Candida albicans*. Curr Opin Microbiol. 2006;9(6):595–601.

4. Sudbery P, Gow N, Berman J. The distinct morphogenic states of *Candida albicans*. Trends Microbiol. 2004;12(7):317–24.

5. Sudbery PE. Growth of *Candida albicans* hyphae. Nat Rev Microbiol. 2011;9(10):737–48.

6. Brown AJ, Brown GD, Netea MG, Gow NA. Metabolism impacts upon Candida immunogenicity and pathogenicity at multiple levels. Trends Microbiol. 2014;22(11):614–22.

7. Erwig LP, Gow NA. Interactions of fungal pathogens with phagocytes. Nat Rev Microbiol. 2016;14(3):163–76.

8. Russell DG, Vanderven BC, Glennie S, Mwandumba H, Heyderman RS. The macrophage marches on its phagosome: dynamic assays of phagosome function. Nat Rev Immunol. 2009;9(8):594–600.

9. Sun-Wada GH, Tabata H, Kawamura N, Aoyama M, Wada Y. Direct recruitment of H^+^-ATPase from lysosomes for phagosomal acidification. J Cell Sci. 2009;122(Pt 14):2504–13.

10. Canton J, Khezri R, Glogauer M, Grinstein S. Contrasting phagosome pH regulation and maturation in human M1 and M2 macrophages. Mol Biol Cell. 2014;25(21):3330–41.

11. Danhof HA, Lorenz MC. The *Candida albicans* ATO gene family promotes neutralization of the macrophage phagolysosome. Infect Immun. 2015;83(11):4416–26.

12. Vylkova S, Carman AJ, Danhof HA, Collette JR, Zhou H, Lorenz MC. The fungal pathogen *Candida albicans* autoinduces hyphal morphogenesis by raising extracellular pH. MBio. 2011;2(3):e00055–11.

13. Vylkova S, Lorenz MC. Modulation of phagosomal pH by *Candida albicans* promotes hyphal morphogenesis and requires Stp2p, a regulator of amino acid transport. PLoS Pathog. 2014;10(3):e1003995.

14. Martínez P, Ljungdahl PO. Divergence of Stp1 and Stp2 transcription factors in *Candida albicans* places virulence factors required for proper nutrient acquisition under amino acid control. Mol Cell Biol. 2005;25(21):9435–46.

15. Silao FGS, Ward M, Ryman K, Wallström A, Brindefalk B, Udekwu K, et al. Mitochondrial proline catabolism activates Ras1/cAMP/PKA-induced filamentation in *Candida albicans*. PLoS Genet. 2019;15(2):e1007976.

16. Ghosh S, Navarathna DH, Roberts DD, Cooper JT, Atkin AL, Petro TM, et al. Arginine-induced germ tube formation in *Candida albicans* is essential for escape from murine macrophage line RAW 264.7. Infect Immun. 2009;77(4):1596–605.

17. Miller SM, Magasanik B. Role of NAD-linked glutamate dehydrogenase in nitrogen metabolism in *Saccharomyces cerevisiae*. J Bacteriol. 1990;172(9):4927–35.

18. Nishimura A, Nasuno R, Takagi H. The proline metabolism intermediate Delta1- pyrroline-5-carboxylate directly inhibits the mitochondrial respiration in budding yeast. FEBS Lett. 2012;586(16):2411–6.

19. Rodaki A, Bohovych IM, Enjalbert B, Young T, Odds FC, Gow NA, et al. Glucose promotes stress resistance in the fungal pathogen *Candida albicans*. Mol Biol Cell. 2009;20(22):4845–55.

20. Balbi HJ. Chloramphenicol: a review. Pediatr Rev. 2004;25(8):284–8.

21. Daugherty JR, Rai R, el Berry HM, Cooper TG. Regulatory circuit for responses of nitrogen catabolic gene expression to the GLN3 and DAL80 proteins and nitrogen catabolite repression in *Saccharomyces cerevisiae*. J Bacteriol. 1993;175(1):64–73.

22. Childers DS, Raziunaite I, Mol Avelar G, Mackie J, Budge S, Stead D, et al. The rewiring of ubiquitination targets in a pathogenic yeast promotes metabolic flexibility, host colonization and virulence. PLoS Pathog. 2016;12(4):e1005566.

23. Sandai D, Yin Z, Selway L, Stead D, Walker J, Leach MD, et al. The evolutionary rewiring of ubiquitination targets has reprogrammed the regulation of carbon assimilation in the pathogenic yeast *Candida albicans*. MBio. 2012;3(6).

24. Han TL, Cannon RD, Gallo SM, Villas-Boas SG. A metabolomic study of the effect of *Candida albicans* glutamate dehydrogenase deletion on growth and morphogenesis. NPJ Biofilms Microbiomes. 2019;5:13.

25. Westman J, Moran G, Mogavero S, Hube B, Grinstein S. *Candida albicans* hyphal expansion causes phagosomal membrane damage and luminal alkalinization. MBio. 2018;9(5).

26. Clemmons AW, Lindsay SA, Wasserman SA. An effector peptide family required for *Drosophila* toll-mediated immunity. PLoS Pathog. 2015;11(4):e1004876.

27. Lorenz MC, Bender JA, Fink GR. Transcriptional response of *Candida albicans* upon internalization by macrophages. Eukaryot Cell. 2004;3(5):1076–87.

28. Tebung WA, Omran RP, Fulton DL, Morschhäuser J, Whiteway M. Put3 positively regulates proline utilization in *Candida albicans*. mSphere. 2017;2(6).

29. Ljungdahl PO, Daignan-Fornier B. Regulation of amino acid, nucleotide, and phosphate metabolism in *Saccharomyces cerevisiae*. Genetics. 2012;190(3):885–929.

30. Palkova Z, Devaux F, Icicova M, Minarikova L, Le Crom S, Jacq C. Ammonia pulses and metabolic oscillations guide yeast colony development. Mol Biol Cell. 2002;13(11):3901–14.

31. Ricicova M, Kucerova H, Vachova L, Palkova Z. Association of putative ammonium exporters Ato with detergent-resistant compartments of plasma membrane during yeast colony development: pH affects Ato1p localisation in patches. Biochim Biophys Acta. 2007;1768(5):1170–8.

32. Fernandes TR, Segorbe D, Prusky D, Di Pietro A. How alkalinization drives fungal pathogenicity. PLoS Pathog. 2017;13(11):e1006621.

33. Liao WL, Ramon AM, Fonzi WA. GLN3 encodes a global regulator of nitrogen metabolism and virulence of *C. albicans*. Fungal Genet Biol. 2008;45(4):514–26.

34. Vyas VK, Barrasa, M.I., Fink, G.R. A C*andida albicans* CRISPR system permits genetic engineering of essential genes and gene families. Sci Adv. 2015;1(3):1–6.

35. Vyas VK, Bushkin GG, Bernstein DA, Getz MA, Sewastianik M, Barrasa MI, et al. New CRISPR mutagenesis strategies reveal variation in repair mechanisms among fungi. mSphere. 2018;3(2).

36. Farboud BMBJ. Dramatic enhancement of genome editing by CRISPR:Cas9 through improved guide RNA design. Genetics. 2015;199:959–71.

37. Reuss O, Vik A, Kolter R, Morschhäuser J. The *SAT1* flipper, an optimized tool for gene disruption in *Candida albicans*. Gene. 2004;341:119–27.

38. Tumusiime S, Zhang C, Overstreet MS, Liu Z. Differential regulation of transcription factors Stp1 and Stp2 in the Ssy1-Ptr3-Ssy5 amino acid sensing pathway. J Biol Chem. 2011;286(6):4620–31.

39. Davis MM, Alvarez FJ, Ryman K, Holm AA, Ljungdahl PO, Engström Y. Wild- type *Drosophila melanogaster* as a model host to analyze nitrogen source dependent virulence of *Candida albicans*. PLoS One. 2011;6(11):e27434.

40. Morales DK, Grahl N, Okegbe C, Dietrich LE, Jacobs NJ, Hogan DA. Control of *Candida albicans* metabolism and biofilm formation by *Pseudomonas aeruginosa* phenazines. MBio. 2013;4(1):e00526–12.

41. Silve S, Volland C, Garnier C, Jund R, Chevallier MR, Haguenauer-Tsapis R. Membrane insertion of uracil permease, a polytopic yeast plasma membrane protein. Mol Cell Biol. 1991;11(2):1114–24.

42. Liu H, Kohler J, Fink GR. Suppression of hyphal formation in *Candida albicans* by mutation of a *STE12* homolog. Science. 1994;266(5191):1723–6.

43. Lee KL, Buckley HR, Campbell CC. An amino acid liquid synthetic medium for the development of mycelial and yeast forms of *Candida albicans*. Sabouraudia. 1975;13(2):148–53.

44. Martínez P, Ljungdahl PO. An ER packaging chaperone determines the amino acid uptake capacity and virulence of *Candida albicans*. Mol Microbiol. 2004;51(2):371–84.

45. Vonk AG, Wieland CW, Netea MG, Kullberg BJ. Phagocytosis and intracellular killing of *Candida albicans* blastoconidia by neutrophils and macrophages: a comparison of different microbiological test systems. J Microbiol Methods. 2002;49(1):55–62.

46. Fonzi WA, Irwin MY. Isogenic strain construction and gene mapping in *Candida albicans*. Genetics. 1993;134(3):717–28.

47. Dabas N, Morschhäuser J. Control of ammonium permease expression and filamentous growth by the GATA transcription factors *GLN3* and *GAT1* in *Candida albicans*. Eukaryot Cell. 2007;6(5):875–88.

